# The Wnt co-receptor Arrow-LRP5/6 is required for Planar Cell Polarity establishment in *Drosophila*

**DOI:** 10.1101/2025.09.22.677847

**Authors:** Ursula Weber, Reza Farhadifar, Marek Mlodzik

## Abstract

Wnt-signaling, via β-catenin or the planar cell polarity (PCP) branch, is crucial for development, tissue homeostasis, and linked to many diseases. LRP5/6, *arrow* (*arr*) in *Drosophila*, is the obligate co-receptor in Wnt/β-catenin signaling, with ligand binding to a Frizzled (Fz) family member and LRP5/6 mediating formation of the signalosome complex with Dishevelled (Dsh/Dvl in mammals) and Axin. Current models for Wnt/PCP signaling omit Arr/LRP5/6 and the notion is that it functions without these co-receptors. Here we show that *arr/LRP5/6* is positively required in Wnt/PCP signaling. In *Drosophila*, loss of *arr* results in PCP mediated cellular orientation defects, aberrant wing hair formation, and loss of polarity, as described for core PCP factors *fz, fmi/Celsr,* and *dsh*. In the eye, *arr* mutant tissue displays cell fate changes in photoreceptors R3/R4 and chirality defects, classical PCP phenotypes. During Wnt/PCP establishment, defects are manifest as reduced levels of Fmi/Celsr and Dsh along with loss of their asymmetric localization. Functional interactions indicate that Fz can recruit Arr, and this potentiates Fz and Dsh function in PCP signaling in all tissues tested. Taken together, our data support an essential Arr/LRP5/6 function in promoting Wnt/Fz-Dsh PCP-complex activity.

## Introduction

The Wnt family of signaling molecules is critical for development and organogenesis across all animal species, including tissue homeostasis through stem cell maintenance and patterning, and linked to many diseases ^1–5^. Wnt signals regulate a complex network of related, but distinct, signaling pathways. The historically most studied Wnt-pathway acts through β-catenin to activate transcription, while the best known β-catenin-independent Wnt signaling cascade regulates cell polarity and tissue organization, generally referred to as Wnt/Planar Cell Polarity (PCP) signaling, with Wnt/Frizzled (Fz in *Drosophila*, or Fzd in mammals) ligand-receptor interactions constituting an evolutionarily conserved upstream activating signaling module for all Wnt-pathways ^1–5^. The Wnt co-receptor LRP5/6, Arrow (*arrow/arr*) in *Drosophila*, is thought to be specific for Wnt/β-catenin signaling, and a key component of the signalosome upon Wnt-ligand binding (reviewed in ^2,6–10^). It is broadly understood that the PCP pathway functions without the LRP5/6 co-receptors, and all current models do not consider Arr/LRP5/5 as an activator of Wnt/PCP signaling ^11–13^. Previous studies have ascribed any polarity-related Arr/LRP5/6 phenotypes to indirect effects of earlier requirements for Wnt/β-catenin signaling ^14,15^. In *Drosophila*, *arr* function has been mainly analyzed in embryonic and wing development ^16,17^, and in setting up the polar-equatorial eye field ^14^. These functions have been shown to involve and align with Wnt/β-catening signaling ^16–20^. In vertebrates LRP5/6 is associated with Wnt/β-catening signaling as the obligate co-receptor acting together with Frizzled family receptors (see review references above). Co-receptor involvement in the Wnt/PCP pathway is more complex ^6^. Besides the critical Frizzled family members acting in Wnt/PCP, studies in vertebrates implicate the Receptor Tyrosine kinase-like orphan receptor 1 and 2 (ROR1 and 2), Protein Tyrosine Kinase 7 ^21^, and RYK (Tyrosine kinase related receptor) as co-receptors for the PCP pathway (reviewed by ^7,22,23^), whereas in *Drosophila* no co-receptor for the core PCP pathway has been described.

Wnt ligands generally cluster Frizzleds and co-receptors, which has been best demonstrated in Wnt/β-catenin signaling for Frizzleds and Arr/LRP5/6 (reviewed by ^2,6,10,24^) and, accordingly, a *fz2::arr* fusion in *Drosophila* can bypass the ligand requirement for Wnt/β-catenin signaling activation and function ^17^. In *Drosophila*, both *fz* and *fz2* can act redundantly as receptors for Wnt/β-catenin signaling ^25–28^. However, Fz2 appears to be the primary receptor for Wnt (Wg)/β-catenin signaling, as each of the two Frizzled family members causes different and specific effects when overexpressed/”activated” ^16,29–32^, with Fz2 inducing specifically Wnt/β-catenin phenotypes, whereas Fz causing exclusively PCP defects. Epistatically, *arr* acts downstream of *wg/Wnts* and upstream of *dishevelled* (Dsh, Dvl in mammals), and importantly a membrane tethered version of Dsh can rescue *arr* loss-of-function embryonic lethality and associated Wg/β-catenin signaling defects ^16,33^.

*Drosophila* has been an exceptional model system in defining PCP establishment mechanisms and identifying the functions of its core components ^11–13,34–37^, with analyses primarily in the wing and eye. In wings, each cell is polarized in the proximal-distal axis and produces an actin-based hair (trichome) on its distal vertex (reviewed in ^38^). In the eye, PCP signaling sets up the cell fates of photoreceptor R3 and R4 subtypes, which orients the ommatidial clusters by defining their arrangement within the retinal epithelium (reviewed in ^35,39–41^). PCP establishment depends on the regulated interactions of the Wnt/PCP core factors, that form polarized trans-membrane intercellular complexes within the proximo-distal cell junctions in wing cells or at the R3-R4 precursor interface in the eye ^42,43^. The protein complexes formed by core PCP proteins are a Fz-Dsh-Dgo (*diego*, also Inversin/Diversin in vertebrates) complex distally in wing cells or on the R3 side of the R3/R4 interface in eye discs. The second complex contains the transmembrane protein Van Gogh (Vang, Vangl1/2 in mammals, known as Stbm in *Drosophila*) and its cytoplasmic partner Prickle (Pk, Pk1-3 in vertebrates). Both complexes are stabilized across cell membranes via the homotypic cell adhesion protein Flamingo (Fmi, Celsr in vertebrates, a.k.a. *starry night/stan* in *Drosophila*), which interacts with both Fz and Vang within the membranes. Loss or gain-of-function of any of these PCP core factors affects their asymmetric, polarized localization and hence disturbs PCP orientation in any tissue analyzed ^35,43^. Highly similar behavior of the Wnt/PCP core factors is apparent in vertebrates, with the addition of redundancy – there are several related proteins present for each of the core factors – and the proposed involvement of co-receptors ^34,36,37^. In addition, the Wnt family members Wnt5a and Wnt11 have been identified genetically in vertebrates (in zebrafish) for a PCP specific requirement ^44,45^. In *Drosophila*, Wg and Wnt4 have been proposed to act redundantly in Wnt/PCP signaling ^46^. Similar to the core factors, loss and localized misexpression of Wnts interfere with PCP orientation ^46^.

A critical gap exists in the understanding of the activation of individual, specific Wnt-pathway branches and their selection, as the Frizzled receptors and Wnts are often not pathway branch specific. With the premise to identify additional factors in the context Wnt-signaling pathway selection that might act near or at the level of the receptor complexes, we used Dsh as a genetic tool, because it is critically involved in both pathway branches and is at the node of pathway specificity from upstream input and towards downstream effectors ^31,47,48^. We thus established a *Drosophila* strain with a mild *dsh* knockdown, displaying both Wnt/β-catenin and PCP defects. Prior to using this *dsh* tool in a genome-wide screen for genetic modifiers, we tested known Wg/Wnt signaling components, which showed *arr/LRP5/6* to shift the *dsh* phenotypic balance from Wnt/β-catenin to Wnt/PCP signaling and vice-versa, when *arr* levels were manipulated. Arr behaved like the Fz receptors and the Wnts in this context. This led us to examine the potential role of *arr* in the well-defined PCP signaling landscape in *Drosophila*.

Here we demonsrate that *arr* loss-of-function clones display classical PCP defects in wing and eye tissue, and that *arr* acts positively in Wnt/PCP signaling. This contradicts current models, which generally present Arr/LRP5/6 as a Wnt/β-catenin pathway specific Wnt co-receptor. It is a specifc defect, as the clonal wing PCP phenotypes are rescued by Arr coexpression, confirming a PCP requirement of Arr/LRP5/6. We show in pupal wings that in *arr* mutant clonal patches the asymmetric Fmi and Dsh localization is lost and/or misoriented, coincident with their protein levels being reduced. These defects lead to wing hair misorientations, reminiscent of *fz, dsh* or *Vang* loss-of-function (LOF) phenotypes. In the eye, the establishment of photoreceptor fates R3 and R4 is disturbed in *arr* clones and the chiral arrangement of ommatidia is affected. Consistently, genetic manipulations place *arr* function downstream or at the same level as *fz* and *dsh*, indicating that all three act closely together in the PCP signaling network. Importantly, Arr appears to promote Fz and Dsh PCP activity, with its increase potentiating both Fz and Dsh overexpression PCP effects and its LOF/knockdown reducing these. *arr* shows PCP specific defects also in combination with *wg*, *Wnt4* and *fz*. Taken together, Arr behaves functionally like a member of the Fz-Dsh complex in PCP signaling and likely links it to Wg/Wnt4.

## Results

### Dsh function in Wnt/β-catenin and Wnt/PCP signaling is sensitive to Arr levels

Dsh (Dvl in mammals) is the key cytoplasmic signaling adapter, which confers both Wnt/β-catenin and Wnt/PCP signaling to further downstream effectors ^31,49–52^. Knockdown of *dsh* during wing development causes Wnt/β-catenin and Wnt/PCP LOF signaling defects, visible in the loss of wing margin tissue and misorientation of cellular actin-based wing hairs (trichomes), respectively (Supplemental Figure S1A, also ^53^). In a pilot screen with most ligands and transmembrane proteins/receptors acting in the two main Wnt-signaling pathways, we discovered that manipulating *arr/LRP5/6* levels did tilt the *dsh* phenotypic balance, enhancing and suppressing the respective phenotypes when their levels were changed, very similar to effects seen with the Wnt ligands and Fz family receptors (Supplemental Figure S1A-C and A’-C’, also Supplemental Table 1). As ligands and receptors are shared between the two pathways and *arr* modified the *dsh* knockdown in the same way, we decided to investigate *arr* function in PCP establishment.

### *arr* is required for correct trichome orientation, a Wnt/PCP read out

A key read-out of PCP signaling in the *Drosophila* wing is the distal positioning of actin polymerization and prehair formation within each wing cell, leading to distally oriented actin based hairs, called trichomes ^38^. This feature is set up and preceded by asymmetric localization of the core PCP factors in the proximal-distal axis ^54,55^. LOF of *fz* and *dsh* cause loss of asymmetric core PCP factor localization and actin polymerization is delayed and initiates in the center of cells, leading subsequently to misoriented trichomes. Importantly, in addition to these cell autonomous effects, non-autonomous prehair misorientations are observed around *fz* and *Vang* (a.k.a. *stbm)* mutant tissue, and induced by Wnt4 misexpression clones ^46,56,57^.

In adult *arr* LOF patches (for both alleles: *arr*^2^, a null allele, and *arr^k^*, an insertion k0813 mutation) we observed wing hair misorientations and formation of multiple trichomes per cell, both hallmark phenotypes of core PCP factor mutations (Figure 1B”, C’) ^38^, and, as has been described before, clones near the margin displayed loss of tissue or notches (Fig. 1B, B’, C, compare to Fig. 1A for *wt* control). *arr*^2^ clones are marked by *shn*, producing thin, kinked wing hairs (Fig. 1B’,B”). Strikingly, we also observed misoriented trichomes in *wild-type* (*wt*) cells next to *shn* marked *arr*^2^ clonal tissue (an example is highlighted in Fig. 1B” with *wt* area marked in blue next to mutant *arr*^2^ tissue, in red, displaying misoriented trichomes). These non-autonomous effects appeared to occur at random positions with respect to the proximal-distal axis to the clone, and the cellular misorientations were seen both towards or away from the mutant tissue (summarized in Fig. 1F). As such, these non-autonomous defects are unique and do not resemble known non-autonomous phenotypes associated with core PCP factors (see above and Discussion). We conclude that *arr* LOF displays autonomous and novel non-autonomous PCP defects in the adult wing.

**Figure 1.**
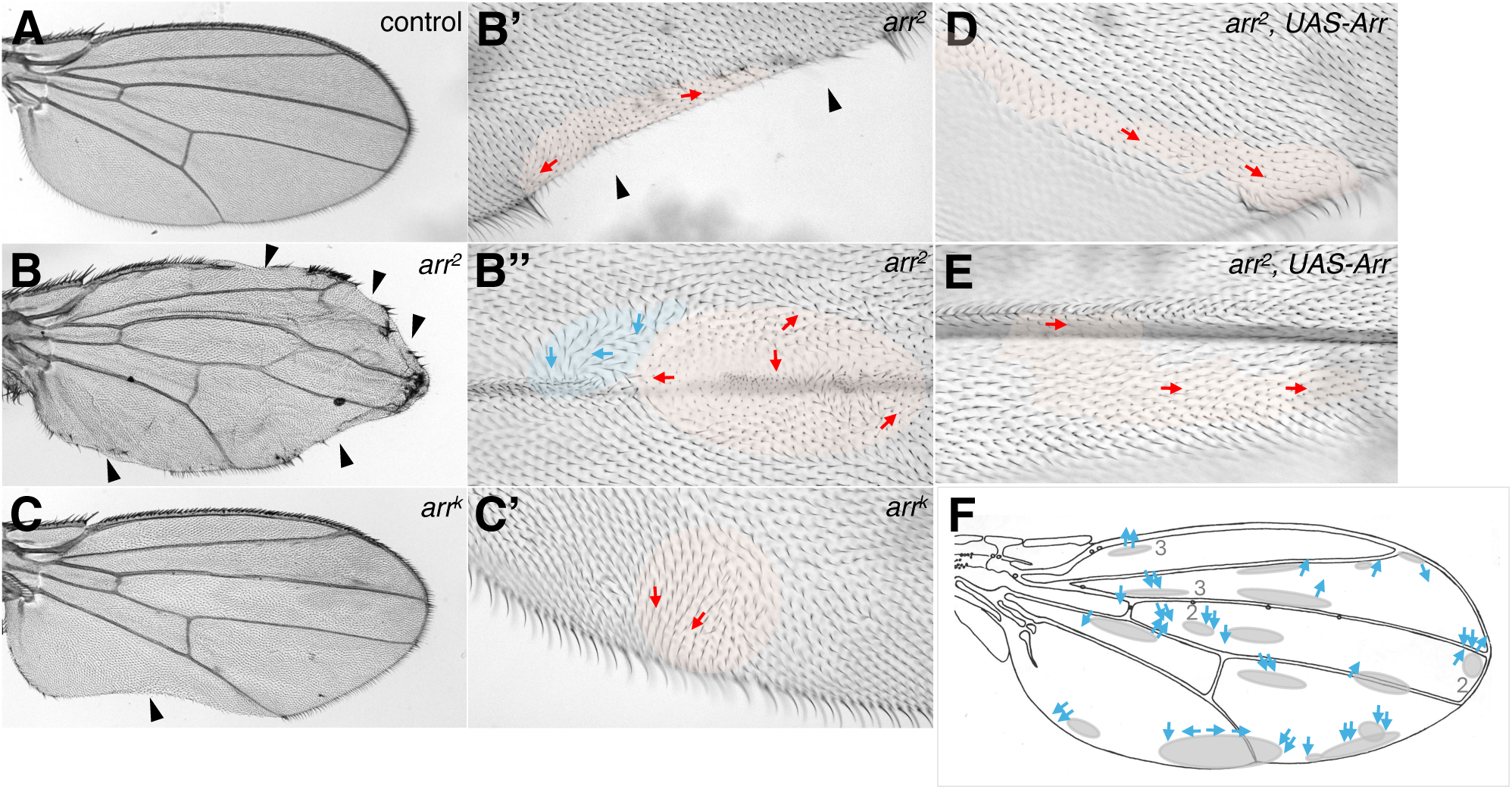
*arr* loss-of-function induces autonomous and non-autonomous PCP defects in the wing. (A-E) Adult wings with clones for *arr*^2^ (B) and *arr^k^* (C), and *arr*^2^ rescue via *UAS-Arr* (D,E). Wing margin loss/notching is indicated by arrowheads in (B,B’,C). (B’) Higher magnification of *arr*^2^ clone, marked by *shn,* induced notching and misoriented trichomes, indicated by red arrows (mutant tissue highlighted in red). Clones for *arr*^2^ (B”) and *arr^k^* (C’) display wing hair misorientation, occasional loss and multiple cellular hairs. *arr*^2^ clones, marked by *shn* (red) non-autonomously affect PCP orientation in neighboring *wild-type* tissue (marked in blue, B”), with misorientation of trichomes indicated by arrows. (F) Summmary of non-autonomous PCP effects associated with *arr*^2^ clones. Wing schematic with clones indicated in grey. Blue arrows mark position and direction of misorientation in adjacent *wt* cells. Most misorientations are parallel to defective orientation just inside mutant clone. Grey numbers indicate how many clones of a similar size were found in a given position. (D,E) *UAS-Arr* expression in *arr*^2^ clones restores wing margin (D) and rescues trichome misorientations (D,E, shown for clones with positions similar to B’, B” for comparison).

To ascertain that *arr* mutations were responsible for these phenotypes we conducted a rescue experiment where LOF clones simultaneously expressed an *arr* rescue construct via the MARCM technique ^58^ (see Methods). Since it has been shown that *arr* dependent embryonic patterning (a Wg/β-catenin phenotype) can be rescued by expression of myristilated-Dsh (myrDsh), we tested such *Dsh*-mediated rescue as well. We observed that Arr, myrDsh and Dsh all rescued the Wg/β-catenin associated loss of wing margin phenotype for *arr*^2^ (Supplemental Figure S1E,F,G,L) and *arr^k^* (Supplemental Figure S1I,J,K,L). Similarly, *arr* knockdown induced notching was rescued by co-expression of Arr (Supplemental Figure S1M-O). In contrast, the Wnt/PCP phenotype, cellular trichome orientation and number, was only rescued by Arr co-expression (Figure 1D,E, Supplemental Figure S1E,I), which is consistent with past observations that Dsh constructs caused severe PCP defects by themselves. As such, expression of Dsh in *arr* clones aggravated the PCP defects of *arr* LOF further. Taken together, these data confirm that both alleles are strong LOF mutations, which can be rescued by Arr co-expression, and that the Wg/β-catenin phenotype in the wing can also be rescued by Dsh overexpression.

### *arr* is required for timely positioning of prehairs and Fmi asymmetric localization

Wnt/PCP signaling in *Drosophila* wings is regulated by the asymmetric localization of the trans-membrane core PCP factors Fmi/Celsr, Vang and Fz, which is critical for cellular orientation in the proximal-distal (P-D) axis ^30,59–61^ at pupal wing stages. The proximal and distal cellular complexes are composed of Vang and Pk, and Fz, Dsh and Dgo, respectively, with Fmi present on both sides providing the intercellular homotypic stabilization ^59,60,62^. Wg/Wnt4 are thought to set up the asymmetric localization of core PCP factors early in pupal development ^46^. The core PCP asymmetry results in the positioning of the wing prehair and its distal orientation, as well as it coordinates cell orientation between neighboring cells in the wing epithelial sheet ^54,55^.

Because *arr* clones were associated with misoriented and multiple trichomes, we next examined developing pupal wings at the time of PCP establishment and actin prehair formation for proximal-distal asymmetric core PCP protein and F-actin localization. Cells mutant for *arr* displayed centrally condensing actin and a delay in forming prehairs, and prehair misorientations (Figure 2A,B). These effects were very reminiscent of *dsh* and *fz* loss-of-function phenotypes ^54,55^. Consistently, Fmi localization in *arr* mutant cells displayed reduced and uncoordinated asymmetry (Figure 2C,C’, and D,D’) as compared to neighboring *wt* control tissue. In contrast, E-cadherin (E-cad), a common component of adherens junctions serving as control, was not affected (Figure 2C”,D”). Fmi asymmetry was evaluated for strength of polarization and direction of asymmetry (nematic order), which is a measure of polarity ^63^ (indicated by red lines in Fig. 2C’,D’ for individual cells). Quantification revealed that mutant cells displayed a loss of or reduced Fmi asymmetry (reduction in length of nematic order lines in Fig. 2C’,D’,E) and had lost their proximal distal coordination as compared to *wt* cells (angle of red lines in Fig. 2C’,D’,F).

**Figure 2.**
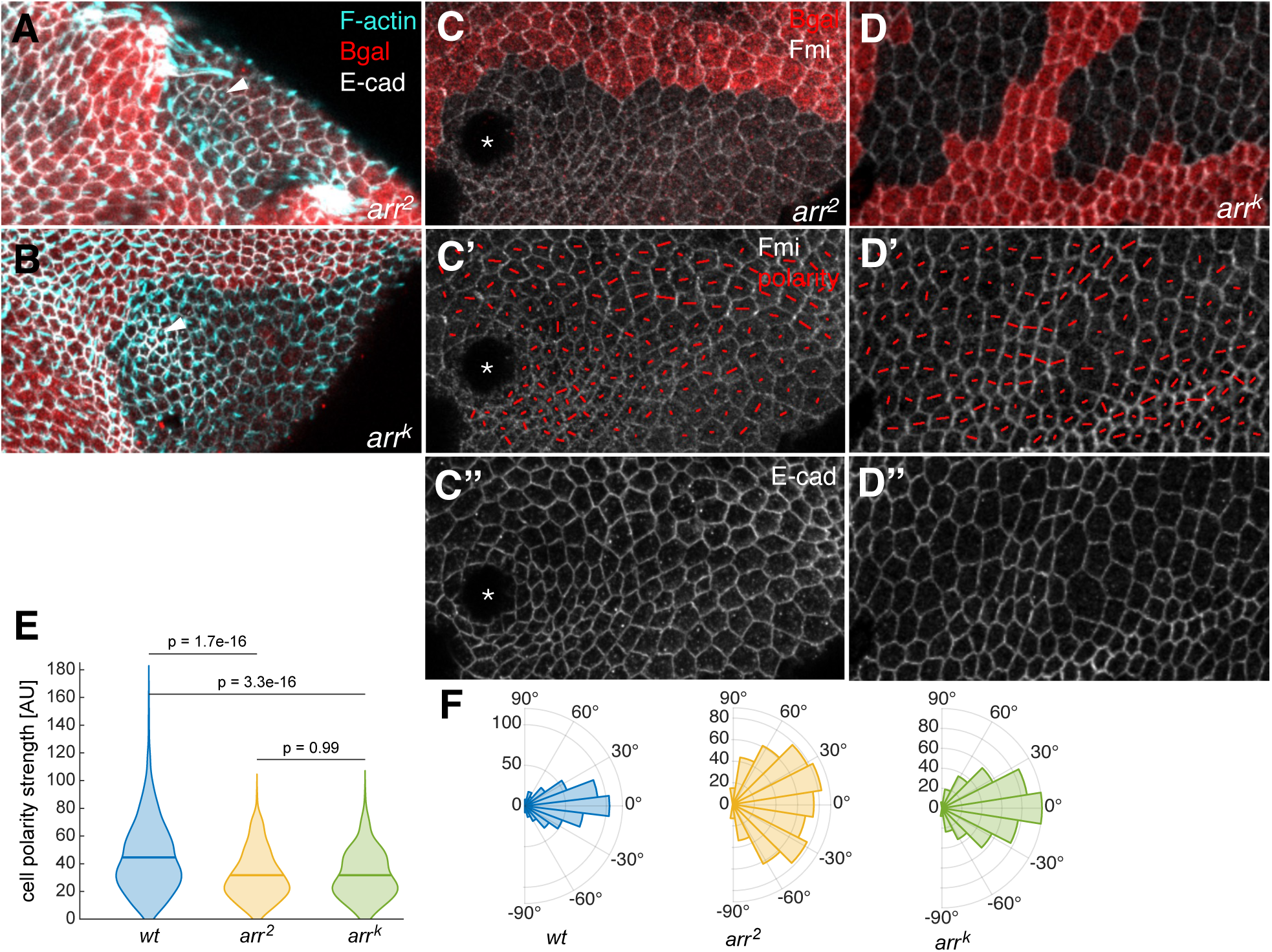
Loss of *arr* delays and mis-positions prehair formation and abolishes Fmi polarization. (A-D) Pupal wings containing *arr* clones. Proximal to the left, anterior at the top. (A,B) Clones of genotype as indicated, marked by the absence of Bgal expression in red. F-actin highlighting prehairs in cyan and E-cadherin (E-cad) for cell outline in white. Note how prehairs form centrally in mutant cells, examples indicated by arrowheads. (C,D) Mosaic tissue stained for Fmi and E-cad. (C’,D’,E,F) Planar cell polarity quantification based on Fmi staining. Polarity (nematic order) strength (length of line) and orientation (angle of line) are indicated by red lines in each cell. Asymmetric Fmi localization is reduced and misoriented in clones compared to *wild-type* tissue (C’,D’). E-cad, as control, is not affected (C”,D”). Tissue outgrowths are observed in some clones, indicated by asterisk (C-C”). (E,F) Quantification of nematic order, polarity strength and direction (angle). (E) Violin plots summarizing polarity strength reduction for both *arr* alleles (C’,D’). (F) Quantification of Fmi asymmetry angles, shown as rosette plots, for *arr*^2^ and *arr^k^* mutant cells. Sectors in rosettes represent average polarity angle distribution. Note that angle distribution is significantly randomized in mutant cells as compared to *wt* control tissue, F-test: *wt* vs *arr*^2^, P=2.655E-07, and *wt* vs *arr^k^*, P=0.0339, from 4-5 wings, 7-8 clones each (see Supplemental Fig. S3 for further comparisons).

Next we examined Dsh, as it is the key scaffold/effector for subcellular downstream effects, including the localization of the actin polymerization machinery ^64,65^, using the DshGFP tool ^52^, which is asymmetrically localized, clustering with Fz and Dgo on distal membranes of pupal wing cells ^60^. This revealed that Dsh levels were markedly reduced and its asymmetric membrane associated localization was lost in *arr* mutant pupal wing cells, before and during prehair formation (Supplemental Fig. S2A,A’,B,B’,C), respectively, whereas E-cad as control was again not affected (Supplemental Fig. S2A”,B”,C).

Taken together, as *arr* LOF affects both Fmi and Dsh asymmetric localization, and coordinated orientation in the proximal-distal axis, and their levels during PCP establishment, we conclude that it plays a key role in PCP signaling, leading to classical PCP defects in pupal wings and manifest in trichome misorientations in adults.

### Arrow is required for photoreceptor R3/R4 fates and PCP establishment in the eye

Precise retinal size and ommatidial arrangement in the eye requires first setting up of the eye field, which includes many inputs as well as inhibition by Wnt/β-catenin signaling ^66,67^, and subsequent cell fate establishment of photoreceptors R3 and R4 via the Wnt/PCP pathway ^68–72^, which ultimately results in a chiral arrangement of photoreceptors in the adult ommatidium, with R3 sitting at the tip and R4 tugged in closer towards the central R7. Notably, two ommatidial chiral forms are established, a ventral and a dorsal, which are organized to form a mirror-symmetric alignment across the dorso-ventral midline, the so-called equator (reviewed in ^41^). This precise arrangement is critical for vision and represents the important role of Wnt/PCP establishment in eye development ^73^.

In order to separate the two Wnt-signaling associated processes of eye field inhibition (Wnt/β-catenin) and cell fate establishment (Wnt/PCP), which in the developing eye are happening in close proximity both temporally and spatially, we first examined adult mosaic *arr* tissue distant from the eye field inhibiting Wg source for cell fate changes via the positioning and chiral arrangement of photoreceptor R3-R4 pairs (Fig. 3A,B, note that both *arr* alleles display very similar defects). Mutant clones near the center of the eye revealed frequent chirality defects, including inversion of the R3/R4 fates or symmetrical cluster arrangements (representative examples are circled in Fig. 3A,B). As Wnt/PCP factors are required in R3, R4 or both ^68–70,74^, we analyzed mosaic ommatidia and R3/R4 pairs in detail for *arr*. Overall, we observed that 13-42% of mosaic R3/R4 pairs had acquired inverted chirality and 10-12% displayed a symmetric R-cell arrangement (loss of chirality) with a double R3-type appearance (summarized in Fig. 1D). The mosaic analysis revealed that there was no bias for which cell of the R3-R4 pair was mutant, to cause these phenotypes, leading us to conclude that *arr* is required in both cells for proper fate establishment (Fig. 3A,B,D). Also, *arr* was not required in or for any other photoreceptor fate. This genetic requirement is similar to that of *fmi* ^70^, among the core PCP factors, and aligns with the pupal wing defects observed with Fmi being reduced and non-polarized in *arr* mutant clones (see also Discussion).

**Figure 3.**
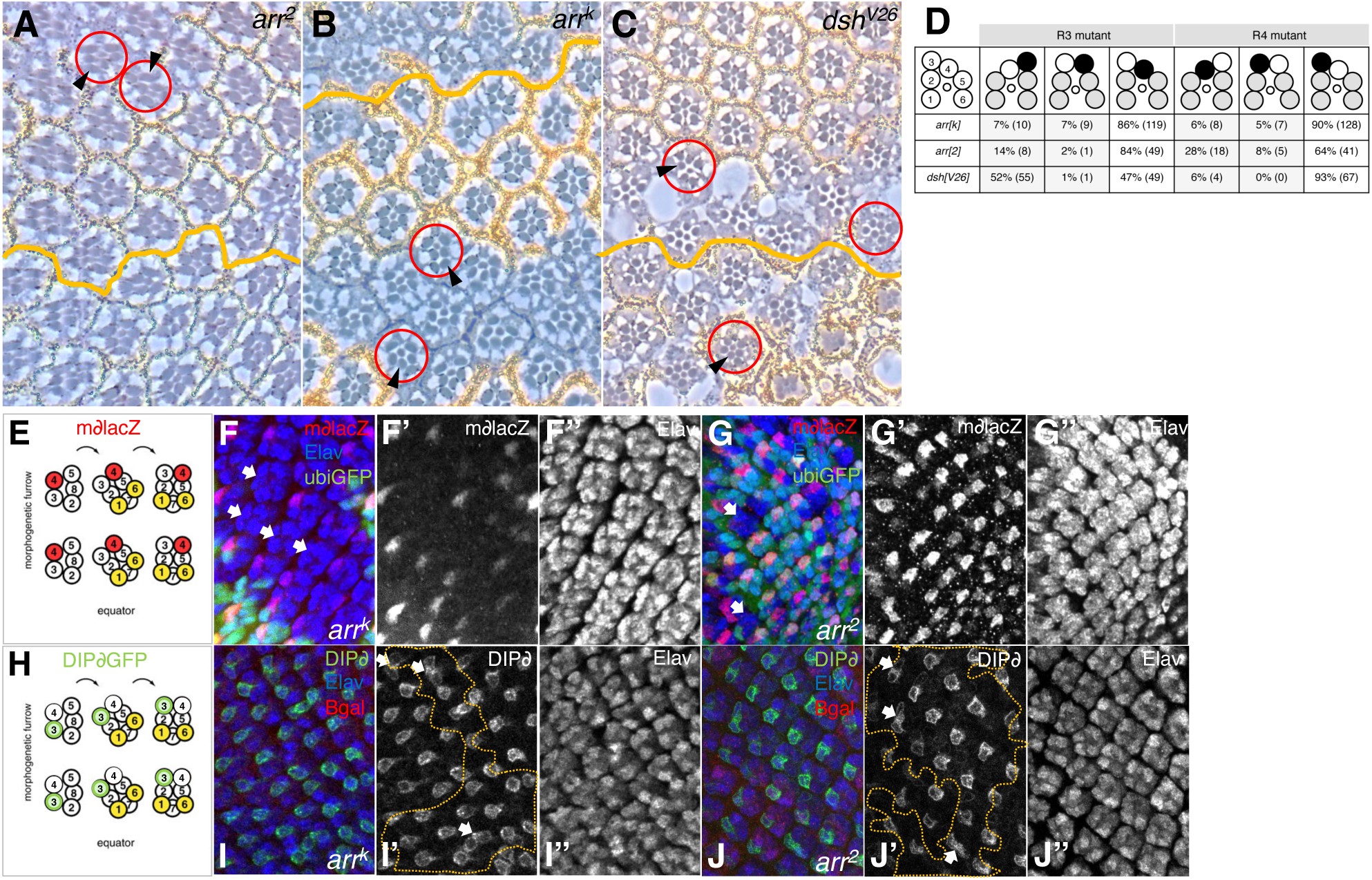
*arr* is required for photoreceptor R3/R4 fate establishment, similar to *dsh*, causing loss of R4 photoreceptor fate. (A-D) Mosaic analysis of photoreceptor pair R3/R4 fates, which are determined by PCP signaling. (A-C) Tangential eye sections of mosaic tissue for *arr* alleles (A,B), and *dsh^V^*^26^ (C) for comparison, as indicated. Anterior is to the left. *Wild-type* tissue is marked by the presence of *white*, seen as yellow pigment granules surrounding each ommatidium and as black dots next to the rhabdomeres in individual R-cells, mutant tissue marked by absence of *white*. (A-C) mutant or mosaic ommatidia display incorrect chirality or PCP defects, circled in red. Black arrowheads point to rhabdomeres of mutant photoreceptors, which have acquired the wrong cell fate and position. Endogenous equator is indicated with an orange line. (D) Summary and quantification of *arr* mosaic analysis, compared to *dsh*. Note *wild-type* rhabdomere (R-cell) arrangement in upper left panel, numbers in circles refer to outer photoreceptor/R-cell identity (small circle represents R7). Panels on right to it show R-cell/rhabdomere arrangement and R3/R4 genotypes. Black circles designate *wild-type* and white circles mutant R3 and R4 cells, with grey filled circles indicating outer R-cell rhabdomeres of *wild-type* or mutant genotype that did not influence ommatidial chirality. Percentage of chirality defects and number of ommatidia (in parenthesis) are shown in lower panels of (D). Grey shaded panels above highlight mutant phenotypic arrangement indicating mosaicism for R3 or R4. Only mosaic ommatidia distant from endogenous eye field inhibiting Wnt/β-catenin source were counted. *arr* mutant R3 or R4 photoreceptors affect chirality establishment in 25-52% of mosaic ommatidia, establishing that *arr* is required in both cells. When *dsh* function is missing in R3, 52% of mosaic ommatidia acquire inverted chirality or a random decision is taken. No effect is seen when R4 is *dsh* mutant, indicating that *dsh* is only required in R3 within the R3/R4 pair. (E-J) Schematics and confocal images of developing ommatidial preclusters in the dorsal half of eye imaginal discs, anterior is left and dorsal up, genotypes and stainings are as indicated. (E) Schematic with photoreceptor R4 fate marker in red (*mδlacZ)*. (F-G) Photoreceptor precursors marked by Elav (blue and monochrome in F”,G”), R4 fate (red, *mδlacZ*, and monochrome in F’,G’) and mutant tissue marked by loss of GFP (green). Arrowheads highlight clusters, which failed to acquire R4 fate (F,G). (H) Schematic with photoreceptor R3 fate marker in green as detected by *DIPδGFP*. (I-J) Photoreceptors are in blue as above, R3 fate in green (*DIPδGFP*) and mutant tissue marked by loss of βgal in red. Mutant tissue is outlined by orange dotted lines. (I’,J’) Arrowheads point to clusters with 2 cells of R3 photoreceptor fate. Note that both *arr* alleles show similar defects in each assay.

As *dsh* is a critical component downstream of Arr and can rescue *arr* LOF effects ^16,33^ (also Fig. 1D,E, and Supplemental Fig. S1C,D,G-I), and *dsh* requirements within the R3/R4 pair have not been established, we included a *dsh* mosaic R3/R4 analysis to compare it to *arr*. Mutant tissue for the *dsh* null allele was again examined in clones distant from the eye-inhibiting Wg source to eliminate potential interference by Wnt/β-catenin signaling (Fig. 3C,D). In *dsh* R3/R4 mosaics, 52% of R3 mutant pairs showed inverted chirality, with mutant R4 mosaics displaying normal chirality (Fig. 3D, 93%), indicating that *dsh* is required in R3 exclusively and its loss results in random chirality decisions. Furthermore, these data indicate that *dsh* is not sufficient to induce R3 fate, like *fz* (ref ^68^), which is required in and sufficient for R3 fate. Taken together, these data suggest that while the *arr* and *dsh* adult eye LOF phenotypes appear similar, their requirements in photoreceptors R3 and R4 differ, indicating additional roles for Arr beyond acting solely through Dsh (see Discussion).

The positions of R-cells in adult eyes are only suggestive of their fate, and we thus examined the R3/R4 precursors in *arr* mutant tissue with specific cell fate markers in developing eye discs. Established tools are for R4 fate *mδ-lacZ* ^71^ and for R3 the newly defined *DIP8GFP* ^75^ (see Fig. 3E,H for representative schematics of ommatidial clusters and the respective R-cell fates). *arr* mutant tissue in developing eyes displayed ommatidial preclusters which had lost *mδ-lacZ* staining, the R4 fate (Fig. 3F,F’,G,G’, note that both alleles behaved similarly). Conversely, we observed double *DIPδGFP* positive clusters in *arr* mutant clones, indicating that 2 cells in a given ommatidium have acquired R3 fate (Fig. 3I,I’,J,J’, note again that both independent *arr* alleles showed same effects). Polar clones were not considered in this study, because they can induce ectopic eye fields and anti-equators due to the Wnt/β-catenin requirement in eye field definition ^66,67^ (see Supplemental Fig. S3). In summary, these analyses confirm what the adult experiments suggested, namely that *arr* is required in both photoreceptors R3 and R4 to establish their cell fates, which is similar to *fmi* function. We conclude that *arr* requirements are different from *dsh* in R3 and R4 fate establishment.

### Arr is apically enriched in pupal wing cells and recruited by Fz overexpression

In *Xenopus*, LRP6 has been shown to localize asymmetricaly in mesodermal cells undergoing convergent extension, a process regulated by PCP signaling ^76^. We therefore attempted to localize Arr via the antibody and its HA-tagged version in pupal wing cells and photoreceptor cells undergoing PCP establishment in *Drosophila*. Arr was shown to localize to punctate structures throughout cells with apical enrichment in imaginal disc cells ^18,77^. We found Arr also to be apically enriched in pupal wing cells at the level of E-cad at the time of PCP establishment (Figure 4A,B). However, no specific pattern in developing photoreceptor cells, nor an asymmetric localization in developing wings, was detected with the available tools, suggesting that either Arr is not asymmetrically localized or that only a small subset of Arr molecules, which might be post-translationally modified, for example phosphorylated, would colocalize with the PCP complexes.

**Figure 4.**
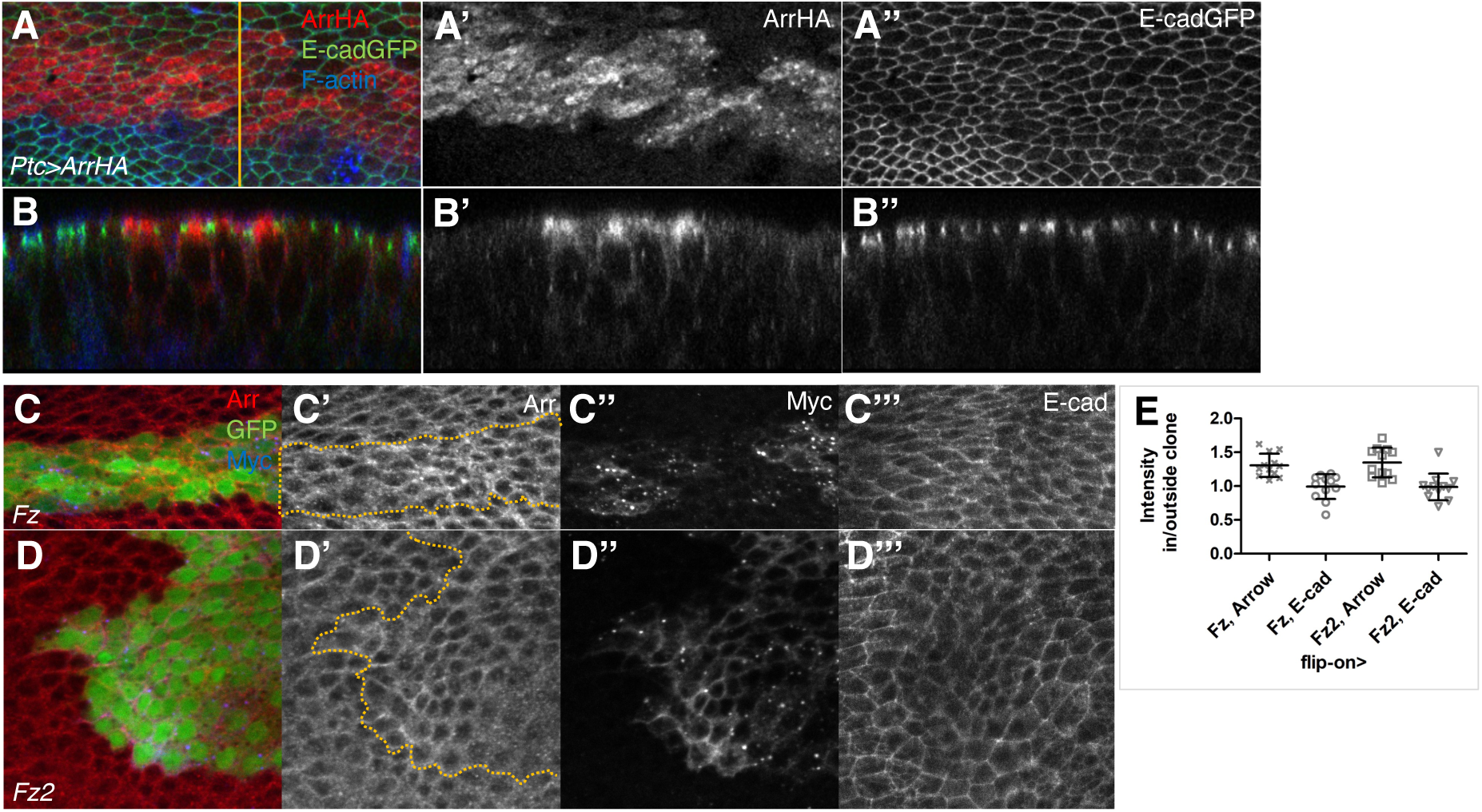
Arr localizes apically in pupal wing cells during PCP establishment and Fz recruits Arr. Confocal images of pupal wing cells expressing Arr-HA via *ptc-GAL4* stained for HA in red, E-cadGFP (green) and F-actin in blue (A,B), proximal is left, anterior to the top (A). Orange line indicates position of Z-line scan images, shown in (B-B”). Arr is strongly enriched in the apical part of wing cells (A’,B’ as illustrated by ArrHA) at the level of E-cad, monochrome in A”,B”. (C,D) Overexpression of Fz increases Arr protein levels. Confocal images of peripodial eye disc cells, anterior to the left, mosaic for either *FzMyc* (C) or *Fz2Myc* (D), with expressing cells marked by GFP, stained for Arr (red, monochrome in C’,D’) and Myc in blue (C”,D”). (C’,D’) Overexpressing clones are outlined with orange line and show punctate peripheral and cytoplasmic elevated Arr levels (in white). (C’’,D’’) Fz and Fz2, tagged by Myc, shown as monochrome. (C”’,D”’) E-cad expression serves as control. (E) Quantification summarizing Arr and E-cad level changes in Fz and Fz2 overexpression clones, as compared to neighboring *wild-type* tissue. E-cad levels are not affected. Arr levels are increased in clones, t-test, P=0.0003 for both *fz* and *fz2* compared to E-cad, from 5-7 discs, 12 clones each.

As *arr* has mostly been studied in Wnt/β-catenin signaling, we wanted to know if Fz GOF, inducing specifically Wnt/PCP signaling, could recruit more Arr to its complexes. To this end we clonally overexpressed Myc-tagged Fz and Fz2 ^32,78^ in developing eye and wing discs and analyzed Arr levels and localization in such clones, compared to adjacent *wt* tissue. As imaginal disc cells are in a pseudostratified columnar arrangement and Arr localization is punctate throughout the apical cytoplasm and periphery of cells ^18,77^, we examined the flat cuboidal cells of the peripodial cell layer of discs. Upon increasing Fz and Fz2 levels clonally we detected elevated Arr levels as compared to adjacent *wt* cells (Fig. 4C-E), whereas control staining was not affected (E-cad, Fig. 4C’’’, D’’’,E). The respective adult wings from these experiments displayed the expected wing phenotypes, with Fz overexpression causing Wnt/PCP specific cell trichome misorientations (Supplemental Fig. S4B,B’, compare to S4A,A’) and Fz2 inducing ectopic margin bristles (Supplemental Fig. S4C,C’), confirming the respective activities of the constructs. We conclude that both Fz (PCP signaling specific) and Fz2 (β-catenin signaling specific) receptors can recruit Arr upon overexpression into their respective signaling complexes.

### Arrow promotes Fz and Dsh function in PCP establishment

As mentioned above, Fz and Fz2 can act redundantly as Wg receptors in the context of β-catenin signaling ^25–28^, whereas Fz is the sole *Drosophila* receptor for Wnt/PCP signaling ^56^. In functional overexpression assays, however, Fz and Fz2 each induce specific signaling effects, with Fz acting strictly in PCP and Fz2 in β-catenin signaling ^16,29,30,32^. Dsh aligns with *arr* in LOF effects ^16,52,53^ but in overexpression it shows mostly PCP and growth defects ^29,31^, whereas Arr overexpression induces the same effects as Fz2 ^16^. Furthermore, a *Fz2::Arr-intra* fusion can activate Wnt/β-catenin signaling independent of ligand activation ^17^.

In order to determine at which level *arr* functions in PCP, we conducted several functional genetic experiments with *fz* and *dsh* combined with Arr (summarized in Supplemental Table S1). In the initial genetic screen assay with *dsh* knockdown (Supplemental Fig. S1A-C), we were surprised to find that altering *arr* levels affected the *dsh* phenotypic balance in both Wnt signaling pathways in a similar manner as the Wnts themselves and the *fz*, *fz2* receptors (see above). We next employed Dsh overexpression in the eye (*sep>Dsh*; see Methods), which caused phenotypes in both pathways (Fig. 5B, compare to 5A for *wt*; quantified in Fig. 5F). *arr* knockdown suppressed *sep>Dsh* for both phenotypes (Fig. 5C,D,F) and overexpression increased both respective effects (Fig. 5E,F). These data are consistent with the notion that Arr promotes/is required for both functions of Dsh, not only Wnt/β-catenin signaling. As a second assay, we used Fz overexpression on the thorax (via *pnrGAL4*), which induced exclusively PCP defects (Fig. 5H, compare to *wt* in 5G; quantified in 5K). Strikingly, the GOF Fz-induced PCP defects were reduced by *arr* knockdown (Fig. 5I,K) and, conversely, enhanced by co-expression of Arr (Fig. 5J,K). Consistently, a Fz overexpression (GOF) PCP phenotype in the eye (*sev>Fz*, Supplemental Fig. S5A) was enhanced by Arr co-overexpression (Supplemental Fig. S5B,C). Taken together, these data indicate that in a PCP-specific signaling context, the levels of Arr potentiate Fz activity, demonstrating a positive requirement for Arr in Fz-mediated PCP establishment, acting in synergy with Fz. They further indicate that Arr can potentiate both *dsh* and *fz* function in PCP establishment in all tissues tested, and *arr* knockdown can suppress *dsh* and *fz* GOF phenotypes in several tissues. These functional assays place *arr* parallel or in the same complex as *fz* and *dsh* in the PCP signaling context.

**Figure 5.**
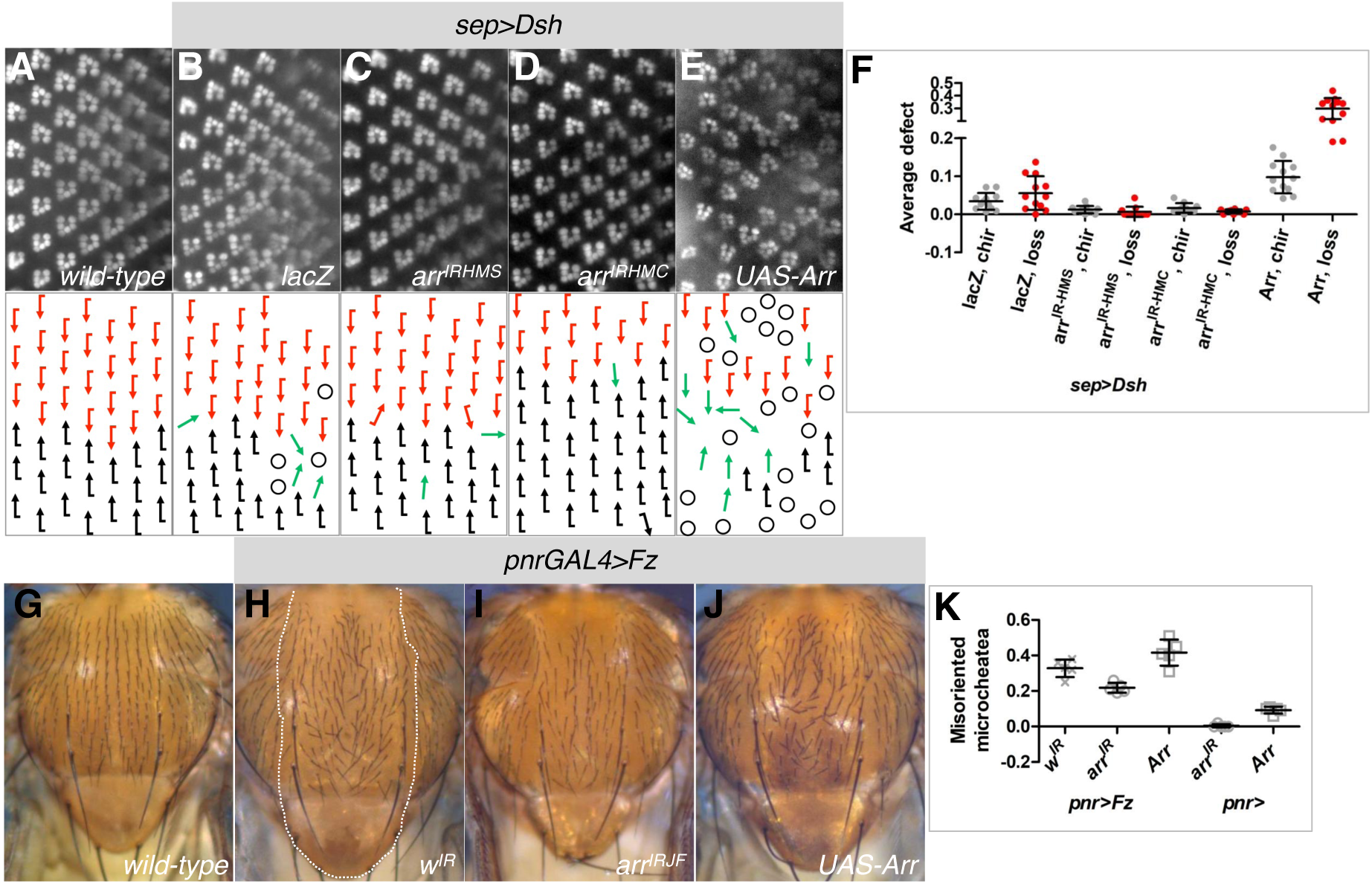
Dsh and Fz induced PCP phenotypes are dependent on Arr levels. (A-E) Ommatidia and their arrangement as seen by fluorescence of the outer photoreceptor rhabdomeres (R1-R6), outlined by *Rh1GFP* in live adult eyes, anterior to the left. In schematics below, red or black arrows represent dorsal or ventral ommatidial chirality, respectively. Green arrows indicate ommatidia with R3-R3 symmetric rhabdomere arrangement and circles outline ommatidia which have lost photoreceptors. Genotypes as indicated with (A) showing wild-type control for comparison. (F) Quantification summarizing genetic interactions. Both phenotypes induced by Dsh overexpression, displaying chirality defects (Wnt/PCP defect, in grey) and loss of photoreceptors (Wnt/βcatenin defect, in red) (B,F), are suppressed by *arr* knock-down (C,D,F) and enhanced by Arr overexpression (E,F). *arr* knockdown or overexpression alone did not induce phenotypes. All interactions are significant compared to *lacZ* controls, t-test: *P*=0.0034, *P*=0.0024, *P*=0.0424, *P*=0.0034, *P*=0.003, *P*<0.0001; 10-12 eyes were evaluated, n=1075-1384 ommatidia each. (G-J) Dorsal notum of *wild-type* (G) and *pnr>Fz* flies (H-J), genotypes as indicated. Microcheatea bristles are misoriented in Fz overexpression area with *pnrGAL4* domain, marked by dotted line in (H). *arr* knockdown reduces (I,K) and overexpression enhances (J,K) Fz effects. (K) Quantification summarizing phenotype in *pnr>Fz* interactions and controls. 5 nota were evaluated each between scutellum, dorso-central bristles and 5 rows anterior to them. 45°–180° misorientations were counted, t-test: *P*=0.0029 and *P*=0.0475 for *arr* knockdown and overexpression, respectively, compared to *w^IR^* control.

To further corroborate the dual role of Arr in Wnt/β-catenin and Wnt/PCP signaling, an *arr* knockdown-induced wing margin loss was rescued by reducing *fz* copy number (Supplemental Fig. S5D,E,G,H) and enhanced by Fz overexpression (Supplemental Fig. S5F,H). These results suggest that Arr is interacting with both receptors, Fz2 and Fz, and the presence or absence of higher levels of the PCP-specifc Fz can titrate Arr away from Wnt/β-catenin to PCP signaling. This is again consistent with the conclusions drawn above that Arr promotes and is required for the PCP activity of Fz and Dsh.

To further test the functional relationships of ligands, receptor (Fz), co-receptor (Arr) and *dsh* in PCP establishment, we generated strains with multiple heterozygous conditions for these components and analyzed their phenotypic effects, if any. A smilar strategy has been used to establish functional links between *wg* and *dsh* ^79^, placing *dsh* in the Wg/β-catenin pathway. In the PCP phenotypic context, a proximal wing area near the anterior crossvein and between veins L3 and L4 has been shown to be sensitive to PCP perturbations (Fig. 6) with, for example, this region displaying a haplo-insufficient defect for *Vang* ^80^ or showing a functional heterozygous interaction between *fz* and *fw* ^81^. We have thus analyzed the cellular trichome orientation in several multiple heterozygous mutant combinations of *fz*, *dsh*, and *arr*, including *wg* and *Wnt4*, the PCP regulating ligands (Fig. 6). Notably, a triple heterozygous genotype of null alleles of *dsh−/+, arr−/+, fz−/+* displayed a 100% penetrance of misoriented cells/trichomes in the proximal wing (Fig. 6B,C), whereas double heterozygous genotypes showed a lower penetrance or no phenotype (Fig. 6A,D; similar but weaker effects were seen with the hypomoprhic *dsh*^1^ allele, see Supplemental Fig. S6 for these and additional allelic combinations). Importantly, heterozygosity in the Wnt family members, *wg and dWnt4*, also increased the penetrance of the PCP defects of *arr−/+, fz−/+* heterozygous animals (Fig. 6C,D; see Supplemental Fig. S6C,D for *wg*−/+ control). Taken together with the above functional interactions (Fig. 5), these experiments demonstrate that *arr* can be rate limiting and thus critical in Fz-Dsh PCP regulation. Moreover, they further support the model that Arr function in PCP is mediated by its interaction with Wg and Wnt4, possibly acting as co-receptor to Fz (see Discussion below).

**Figure 6.**
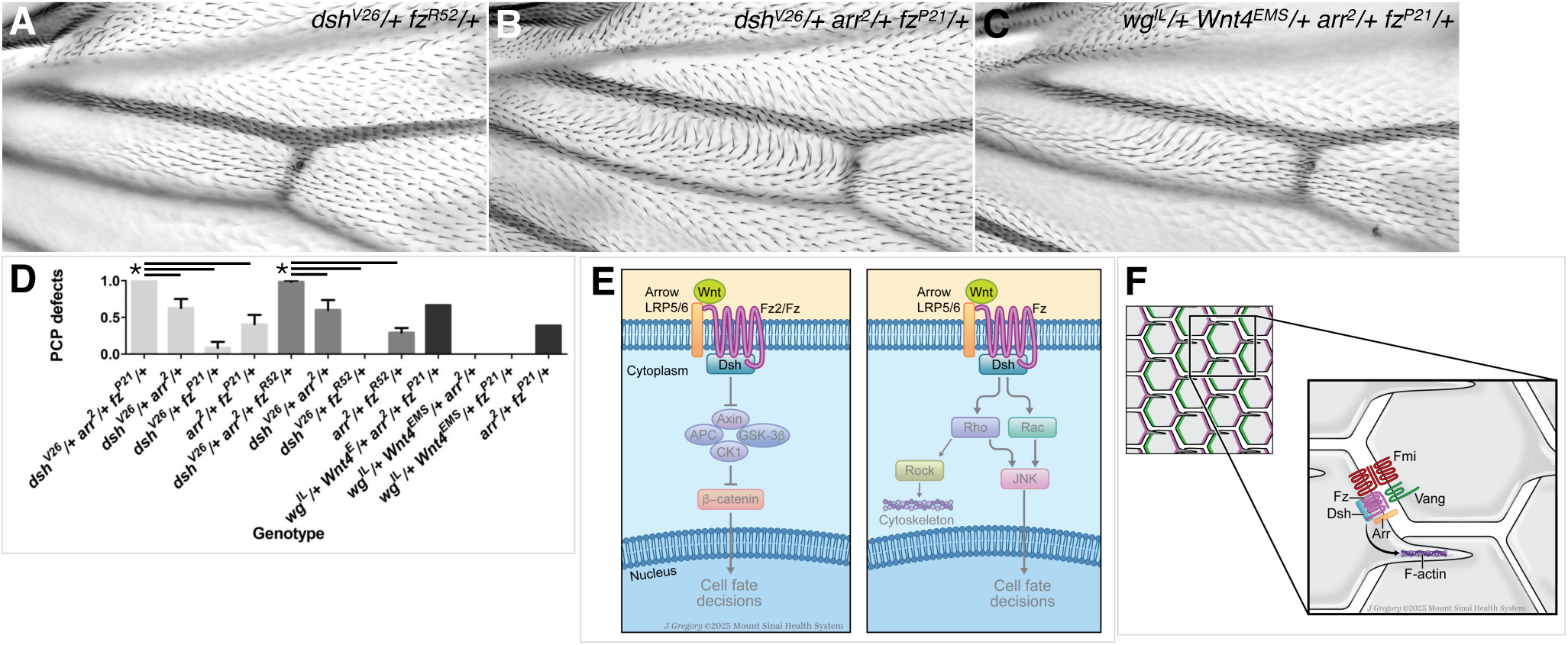
Heterozygosity for *arr*, *fz, dsh* or *wg, Wnt4* result in PCP defects. (A-C) Wings of multiply heterozygous flies for *arr*, *fz*, *dsh* or *wg*, *Wnt4*, with genotypes as indicated. A proximal area, anterior to the anterior crossvein between veins L3 and L4 is shown; this area is sensitive to PCP perturbations (see main text). Examples with significantly increased penetrance of trichome misorientations in *dsh−/+, arr−/+, fz−/+* (B,D), as compared to control double heterozygous *dsh−/+, fz−/+* wings (A,D) are shown, including heterozygosity for ligands *wg* and *Wnt4* in the *arr−/+, fz−/+* context (C,D). (D) Quantification summarizing these interactions shown as % of wings with PCP defects, like examples shown in (B,C). Triple heterozygous *dsh−/+, arr−/+, fz−/+* wings show 100% PCP defect (B,D), compared to *dsh−/+, arr−/+* (62.8%) or *arr−/+, fz−/+* (6.35%) and *dsh−/+, fz−/+* (40.5%). Similar significant effects were found for an independent *fz* allele (97.9%, compared 63.3% or 30.5% and 0%). t-test: *P*<0.001 for triple heterozygotes compared to all double heterozygote controls, n=15-78 wings each. Quadruple heterozygotes *wg−/+*, *Wnt4−/+*, *arr−/+*, *fz-*/+ show 66.7% PCP defects, as compared to *arr−/+*, *fz-*/+ at 38.9% or other double heterozygotes at 0%. t-test: *P*<0.001 for quadruple heterozygotes compared to double heterozygote controls, n=15-18 wings (D). (E,F) Models summarizing the role of Arr/LRP5/6 in PCP establishment. (E) Arr function in Wnt/PCP signaling: Arrow might associate closely with Wnt ligands, the Fz receptor and Dsh and promote their function in PCP (right panel), similar to its role described for the signalosome in Wnt/β-catenin signaling (left panel). (F) Arr acts positively on Fmi asymmetric localization with an effect on F-actin condensation and its distal localization in pupal wing cells. It shows the same requirement as *fmi* in setting up photoreceptor R3 and R4 fate in PCP establishment in the eye.

## Discussion

Here we demonstrate an unexpected role for the Wnt/β-catenin pathway co-receptor Arrow/LRP5/6 in Wnt/PCP signaling, using the *Drosophila* model. Loss-of-function studies with several *arr* alleles reveal classical PCP phenotypes in both wing and eye tissues. Functional mechanistic studies demonstrate that Arr acts in close connection with the Fz/Dsh PCP complex, promoting its activity. Detailed analyses of PCP defects in *arr* LOF pupal wing clones show a loss of polarity, as detected by markedly reduced levels of asymmetric Fmi and Dsh localization, leading to delayed and central formation of the actin-based cellular hairs (trichomes). These features all resemble *fz* and *dsh* LOF PCP phenotypes. Notably, the effects of changing levels of Arr in functional assays with Fz and Dsh indicate that Arr promotes the activity of the Fz/Dsh PCP complex. Furthermore, the heterozygosity assays for *arr*, *dsh* and *fz* support this model, and the same assay for *wg*, *dWnt4*, and *arr* and *fz* supports the notion that the Wg, dWnt4 ligands are critical. Taken together, our data are consistent with the conclusion that Arr/LRP5/6 acts positively in the Fz/Dsh-PCP complex in a Wg/dWnt4 associated manner, and in many ways but not all (see below), behaves like a core PCP factor.

### A role of Arrow/LRP5/6 in PCP signaling

In *Drosophila*, previous studies of *arr* have focused on its role in Wnt/β-catenin signaling, mainly in the embryo and in the context of Wg distribution and gradient formation in wing imaginal discs ^16–18,82^. A past study in eye development ^14^ is also consistent with a function in Wnt/β-catenin signaling with setting up the eye field, but it has not been extended beyond that. Our data here provide clear evidence for a PCP requirement in wing and eye development. Besides the expected defects linked to the Wg/β-catenin pathway in both tissues, the *arr* LOF phenotypes observed are associated with classical PCP features. In pupal wings, the clonal LOF *arr* PCP phenotype resembles the cell-autonomous defects of *fz* and *dsh* ^54,55^, with central and delayed trichome formation, and loss of polarity as determined by Fmi localization. The effects on Fmi localization in pupal wing cells resemble those observed in null mutants of the core PCP factors *fz* and *Vang* ^55^, again reflecting a loss of polarity at this stage. A feature of all PCP factors is that in *wt* pupal wing cells, they display asymmetric membrane-associated localization in the proximodistal axis, with distal localization of Fz-Dsh-Dgo and a proximal Vang-Pk complex, and Fmi co-localizing with both complexes, generating intercellular adhesion to stabilize them across cell membranes. We do not detect asymmetric Arr in pupal wing or eye disc cells, but rather a uniform punctate localization throughout the cell (also ^18^). Expression of HA-tagged Arr ^77^ displays an apical enrichment in disc cells. In pupal wing cells undergoing PCP establishment, we detected ArrHA enriched within the apical plane of the adhesion junctions. In this domain it partially co-localizes with the core PCP factors that are also present in this plane. However, while Arr does not appear to display an asymmetry with the available reagents, it is possible that more sensitive tools or detection of a post-translationaly modified Arr, for example carrying a specific phosphorylation, is enriched with the PCP complexes. Similar issues with CK1ε have initially suggested that it is not asymetrcially localized ^83,84^, which was subsequently revised with higher sensitivity tools establishing that CK1ε is enriched at Fz-Dsh PCP complexes ^85^.

During eye development PCP signaling is key to the R3/R4-pair cell fates, creating a distinction between R3 and R4, which in turn leads to the establishment of ommatidial chirality and giving direction to ommatidial rotation in developing eye tissue (reviewed in ^35,39–41^). Loss of *arr* eye PCP phenotypes have to be analyzed near the equator of the eye field, the dorso-ventral midline, as closer to the poles Wnt/β-catenin signaling defects cause ectopic eye fields and general eye disorganization ^66,67^ (also Supplemental Fig. S3A-E). And here we show that *arr* LOF eye clones also display PCP defects with randomized chirality or clusters remaining symmetrical, again features seen in all core PCP factor mutants. As determined in eye imaginal discs with photoreceptor R3/R4 specific markers, the most striking defect is the frequent loss of the R4 fate and the appearance of R3/R3-type symmetrical preclusters.

### Distinct genetic requirements of *fz*, *dsh* and *arr* in photoreceptor R3/R4 specification

Mosaic analyses within the R3/R4 pair provide additional insight by asking in which cells of the pair genes are required and how they affect the respective cell fate. For example, *fz* and *dgo* are required in R3 to specify R3, while *Vang* and *pk* are required in R4 to specify R4 fate ^68,69,74^. In contrast, *fmi* is required in both cells, as it is thought to provide the adhesive bridge between R3 and R4 stabilizing the Fz and Vang-associated complexes ^70^. Our mosaic analyses of *dsh* and *arr* reveal a distinct requirement from *fz*. While *dsh* is – like *fz* – required in R3, it is not sufficient to push the *wt* cell within the R3/R4 pair towards the R3 fate. Strikingly, *arr* is genetically required in both cells, R3 and R4, resembling the *fmi* requirement. Thus, while in functional interactions during PCP establishment of wing, eye, and thorax Arr promotes Fz and Dsh activity, and the autonomous *arr* LOF phenotypes in the wing largely resemble *dsh* phenotypes, surprisingly *fz*, *dsh*, and *arr* display distinct genetic requirements in R3/R4 specification, and in particular *arr* is required in both cells. PCP establishment in the eye is more complex than in the wing, as it includes here critically the activation of Notch-signaling in R4 via ligand expression in R3 ^71,72^. Dsh has also been suggested to inhibit Notch-signaling, and as such both Dsh and Arr could have effects in both cells, distinct from Fz, preventing Dsh from simply pushing for R3 fate (as seen with Fz) and Arr is possibly required in both cells via Wnt sequestration (see below). Of importance, Notch signaling in R4 upregulates *fmi* expression ^70^, which is itself needed for PCP establishment, with a strict requirement in and sufficiency for R4. As such, Arr and Dsh might contribute directly or indirectly to PCP effects associated with both pathways, Fz/PCP and Notch signaling. Nevertheless, in wing development the cell-autonomous *arr* LOF phenotypes resemble *dsh* phenotypes ^53^ both, for Wnt/β-catenin signaling and the PCP pathway (this study).

### Non-autonomous PCP requirement of *arr*

In addition to the cell-autonomous defects, resembling *dsh*, many *arr* clones also display non-autonomous effects in neighboring *wt* cells, causing their misorientation. This non-autonomy appears, however, different from either *fz* or *Vang* clones ^57,86^, which show non-autonomous effects on the distal side (with misorientation towards the clone) in the case of *fz* or the proximal side (misorientation away from the clone) with *Vang*. Also, they do not mimic the effect of Wnt4 misexpression clones, which reorient cells away from the clone, perpendicular to the clonal border ^46^. Generally, *arr* LOF clones cause a random non-autonomous misorientation of *wild-type* cells, which can be positioned distally, proximally, or even laterally. Nevertheless, the non-autonomous effects often appear to share one feature, with *wild-type* cells tending to take on the same misorientation as adjacent cells within the clone.

What could be causing these non-autonomous defects? A first possibility could arise from the fact that Wnt/β-catenin signaling transcriptionally regulates the graded expression of *four-jointed* (*fj*) and *dachsous* (*ds*), two key components of the Fat/Ds PCP system (reviewed in ^87^). However, an effect through the Fat/Ds system can be excluded, because the respective cell autonomous and non-autonomous PCP phenotypes of *arr* do not resemble those of any of the Fat/Ds component mutants (reviewed in ^87^). Moreover, the cell autonomous phenotypes seen in *arr* wing clones closely resemble core PCP factor mutations, with a loss of polarity as seen with Fmi localization, whereas *fat* or *ds* mutant clones show an upregulation of Fmi and its mispolarization with an increase in actual polarity (reviewed in ^87^). Taken together, our data and these observations strongly argue against effects of Arr via the Fat/Ds system. Furthermore, the non-autonomous effect should not be caused by the reduction in Fmi levels, as seen in *arr* mutant clones, because *fmi* LOF clones do not display non-autonomous defects ^55^.

A potential cause for the non-autonomous cellular misorientations of *arr* could be its effect on Wg/Wnt protein distribution. It has been shown that *arr* loss affects the Wg protein gradient, causing extracellular Wg accumulation in clones ^18^, and as such Arr is critical in shaping the Wg/Wnt protein gradient, in turn affecting the Wnt signaling landscape. In this context, Arr has also been shown to cooperate with Fz2 to degrade Wg in disc cells ^19^, again consistent with shaping Wg distribution in the wing field. The distribution of Wg, and other Wnts, might thus be required for PCP establishment as has been proposed ^46^. This notion can also be reconciled with the observation that *arr* is required in both R3/R4 cells, and its clonal absence could affect the Wg/Wnt levels in the eye as well. Alternatively, Arr might be required to capture Wg/Wnt proteins with Fz to allow Fz-Dsh PCP complex formation and function. In support of this model, overexpression of Fz and Fz2 both sequester increased levels of Arr in co-localizing puncta and compete for Arr association and function (Figures 4-5 and Suppl Fig. S4-5). Both these scenarios strongly suggest that Wnts (Wg/Wnt4) are required for PCP establishment in *Drosophila*, as was demonstrated earlier ^46^. While a Wnt requirement for PCP establishment in *Drosophila* wings has been questioned ^88,89^, these studies provided negative results, with the respective mutant backgrounds displaying significant wing growth, as compared to Wnt knockdowns with focus on Wnt-requirements for wing growth, showing a much reduced wing size ^90^. These comparisons suggest that significant Wnt activity was still present in the respective genotypes ^88,89^ and thus it might not be suprising that little effect on PCP was seen. As Arrow/LRP5/6 is defined as the Wnt co-receptor, the phenotypes reported here add support to a critical role of Wnt proteins in PCP establishment. This conclusion is further supported by the fact that *wg, wnt4* heterozygosity displays a robust PCP phenotype in combination with an *arr−/+, fz−/+* heterozygous condition, suggesting a Fz-Arr receptor/co-receptor function for Wnts in PCP as well (see model in Fig 6E,F),

### Can Arrow/LRP5/6 be considered a core PCP factor?

The data presented here define a new role for the Wnt co-receptor Arr/LRP5/6 in Wnt/PCP signaling and implicate it as a positively acting factor, promoting the function of the Fz-Dsh PCP complex. In that sense it behaves functionally similar to Dgo, which tightly co-localizes with Fz and Dsh in the PCP complex. While Arr functionally interacts with both Fz and Dsh, it does not appear to share the asymmetric, PCP specific, localization. It is largely detectable as punctae in *wt* cells, and, when overexpressed, it is detected in the entire apical junctional cellular region, albeit not in a specific pattern (Fig. 4A,B). Moreover, while overexpression of Fz and Dsh causes strong and specific PCP GOF phenotypes, for example in the eye these genotypes promote R3-R3 symmetrical clusters (e.g. refs^31,72^, Fig. 5B and Supl. Fig. S5A), overexpression of Arr does not generate detectable PCP defects, suggesting that it cannot drive PCP establishment on its own. As such, it is in many ways displaying a unique behavior: while it is required and causing classical PCP phenotypes in LOF assays, it neither displays GOF PCP defects nor does it show a PCP-specific asymmetric localization.

Along these lines, studying LRP5/6 in a PCP context in vertebrates is more difficult due to potential redundancy and also the increased complexity of PCP signaling and associated phenotypes. The existing literature on this topic is limited and, thus far, has possibly suggested an “inhibitory” role. It has been shown in *Xenopus* that LRP6 affects convergent extension, a Wnt/PCP regulated process ^76^, where it has been proposed to antagonize the effect of Wnt11, possibly inhibiting Wnt/PCP signaling. However, this was seen in both LOF and GOF assays, which makes it difficult to interpret. Interestingly, in this study LRP6 is asymmetrically localized in mesodermal cells, on their anterior side, as they undergo the convergent extension process. A second study with focus on convergent extension in vertebrates ^15^ shows that in the mouse *LRP6* LOF phenotypes can be rescued by a deletion of Wnt5a (a presumed PCP specific Wnt). Similarly, an *LRP5/6* knockdown in Xenopus, causing convergent extension phenotypes, is rescued by Wnt5a and Wnt11 knockdowns (both thought to be PCP specific Wnts) ^15^. In contrast to our data presented here, demonstrating a positive Arr/LRP5/6 requirement in PCP establishment by promoting Fz and Dsh PCP function, these convergent extension studies suggest that LRP5/6 might act antagonistically to Wnt/PCP signaling.

In summary, our data demonstrates that in *Drosophila* Arrow/LRP5/6 is a positively acting factor in Wnt/PCP establishment, functionally intertacting with and promoting the cell-autonomous PCP activities of Fz and Dsh. While in the LOF assays it is behaving similarly to core PCP factors, it does not appear to tightly co-localize with these or display a PCP overexpression phenotype on its own. Its non-autonomous PCP requirement, affecting *wild-type* cells adjacent to mutant *arr* LOF clones, could be caused by its effects on Wnt protein distribution.

## Acknowledgments

We are grateful to the Bloomington stock center, Gary Struhl, Nicholas Tolwinski, Jessica Treisman, Nick Baker, Patricio Olguin, Steve Cohen, Konrad Basler, Jeffrey Axelrod, Paul Adler, Sarah Bray, Jun Wu, and Claude Desplan for *Drosophila* strains, to Tadashi Uemura for mouse-αFmi and to DSHB for antibodies. We thank Chaamy Yappa for technical support and Jill Gregory for preparing the model figures. We are also grateful to Alison May, Rob Kraus, Nicholas Tolwinski, and Jun Wu for critically reading the manuscript, and all lab members for support throughout this work. Confocal microscopy was performed at the ISMMS Microscopy CoRE Facility, which was in part supported by the Tisch Cancer Institute P30 CA196521 grant from the NCI. This work was supported by the National Institute of Health (NIH)/NIGMS grant R35 GM127103 to MM.

## Author contributions

U.W. and M.M designed the experiments and wrote the manuscript. U.W. performed the experiments and analyzed all data with help from R.F. who extracted and analyze the polarity associated nematic order data.

## Methods

### Drosophila strains

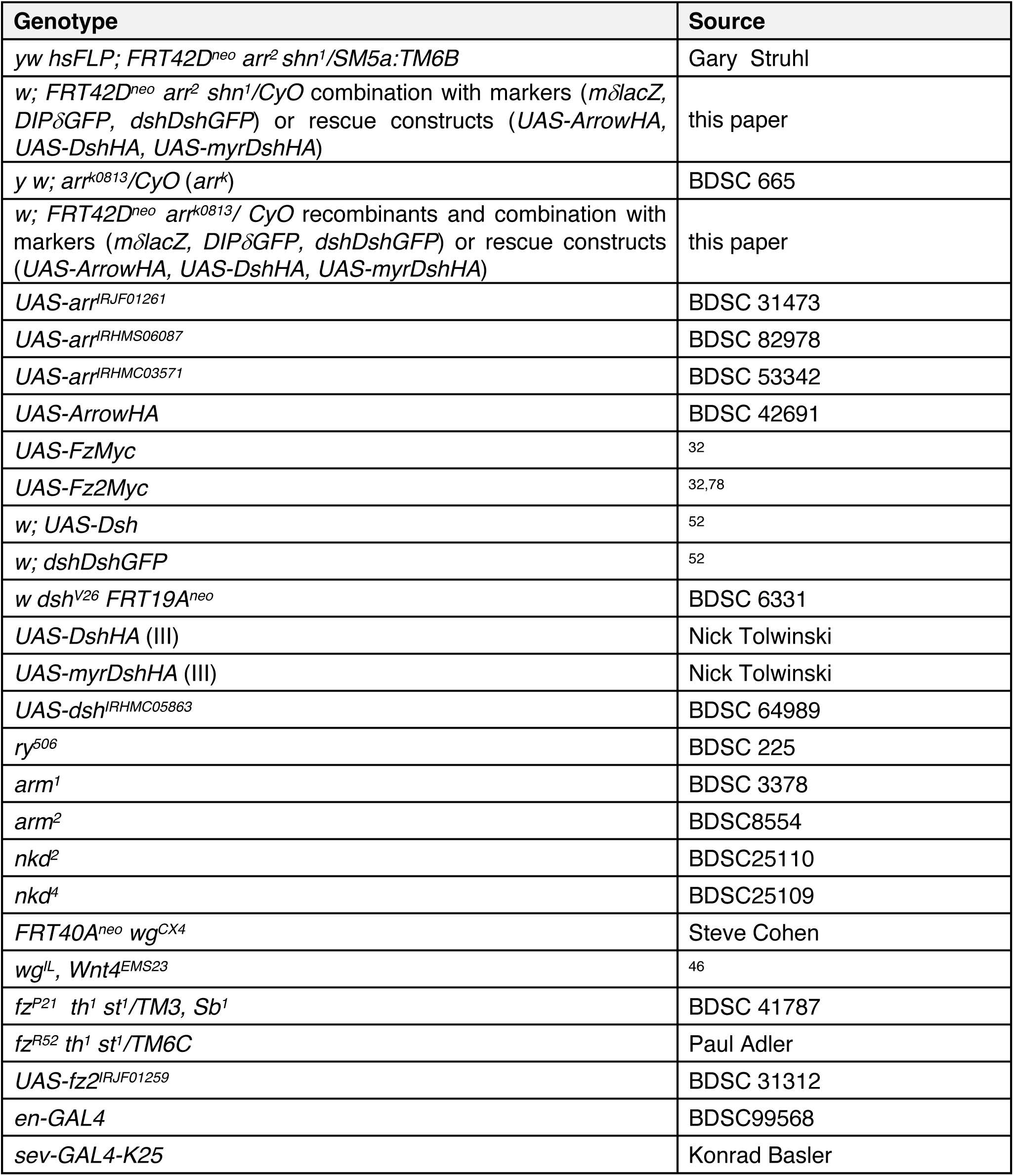
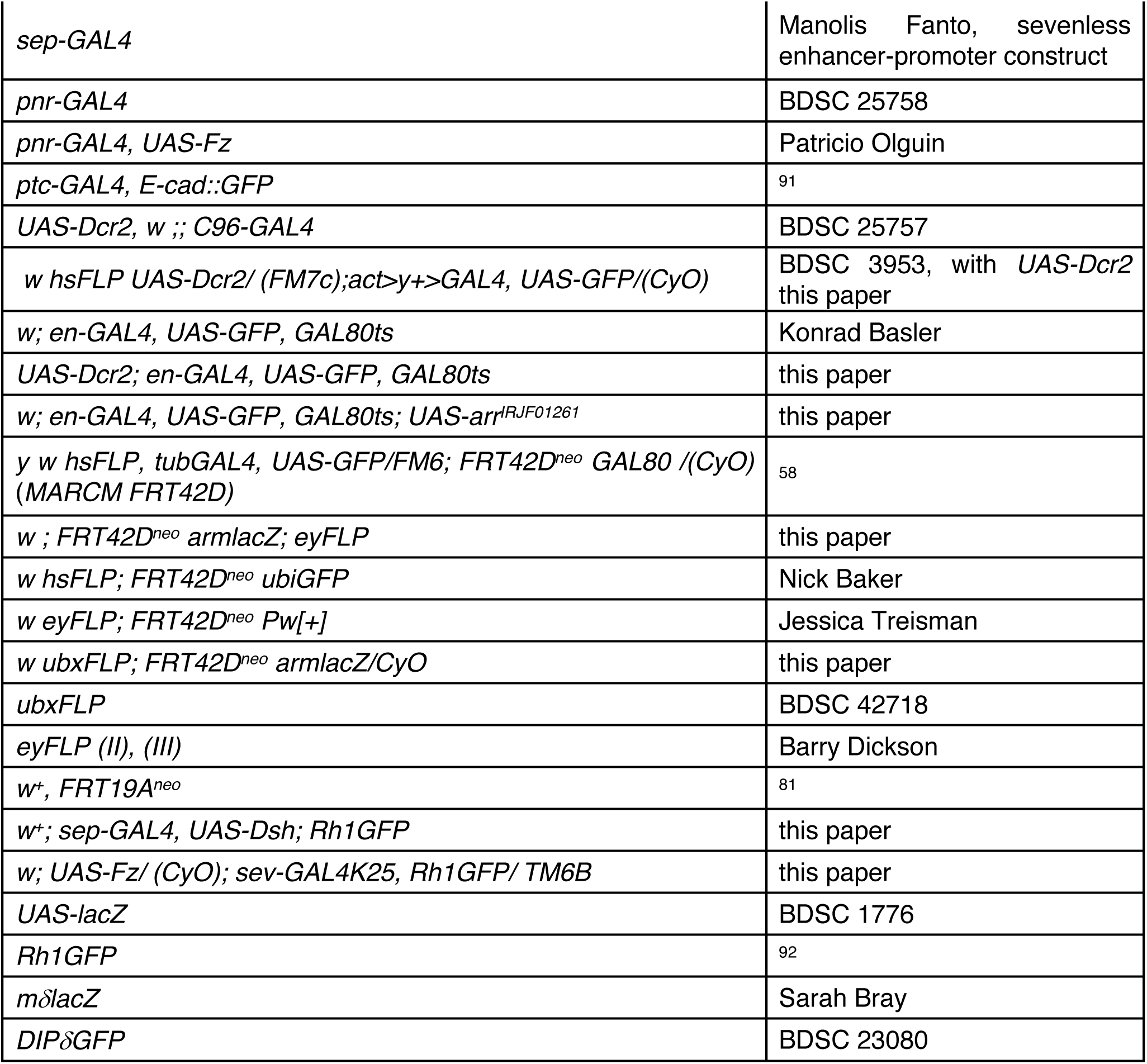

### Genotypes and experimental details by figure

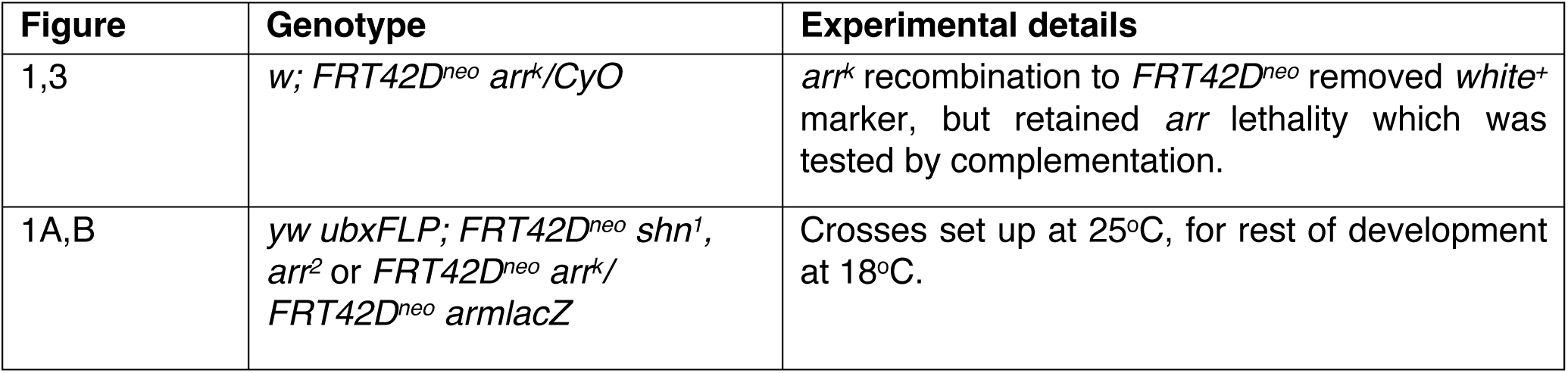
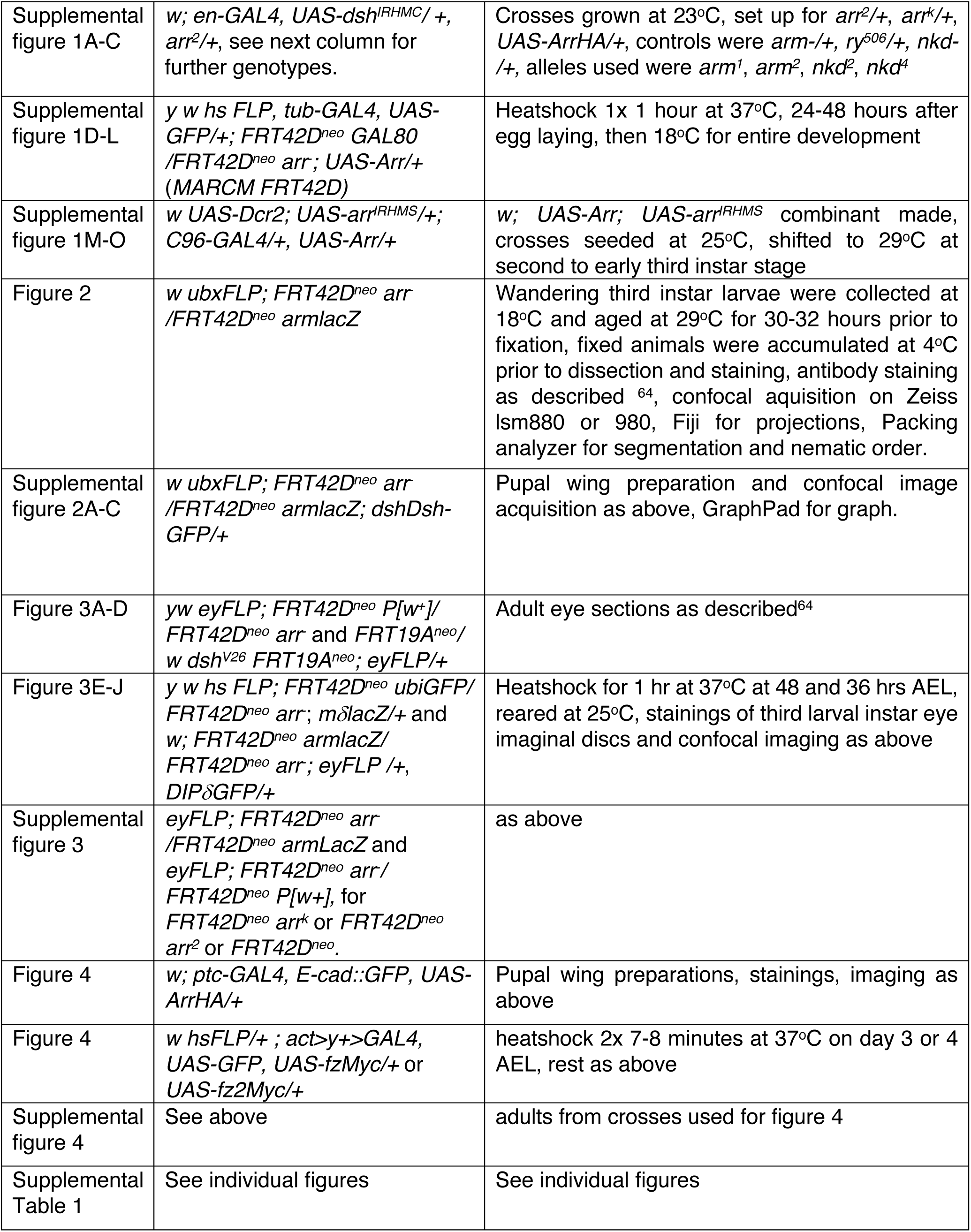
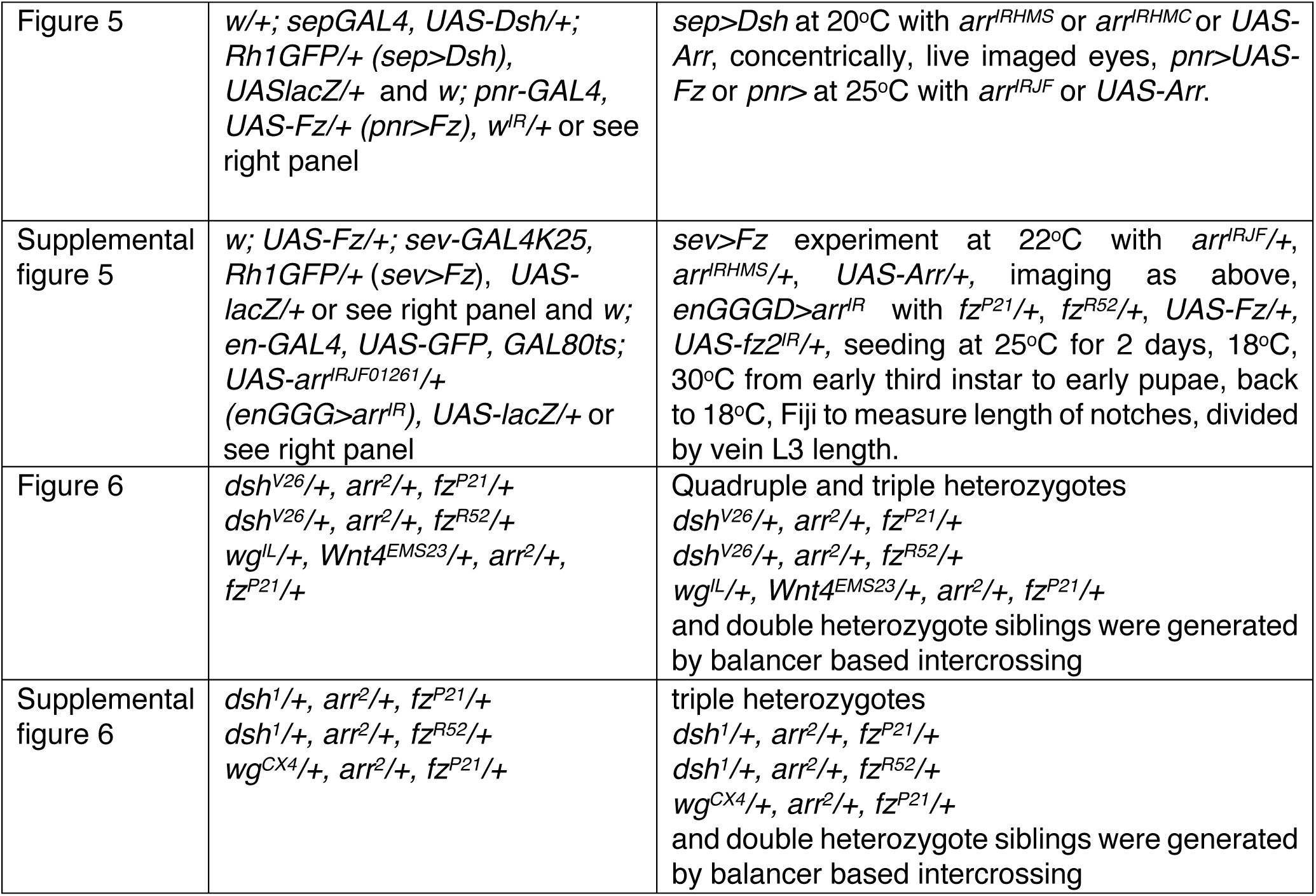

### Immunostaining and histology

Antibody stainings and adult eye embedding as described before ^64^.

### Antibodies

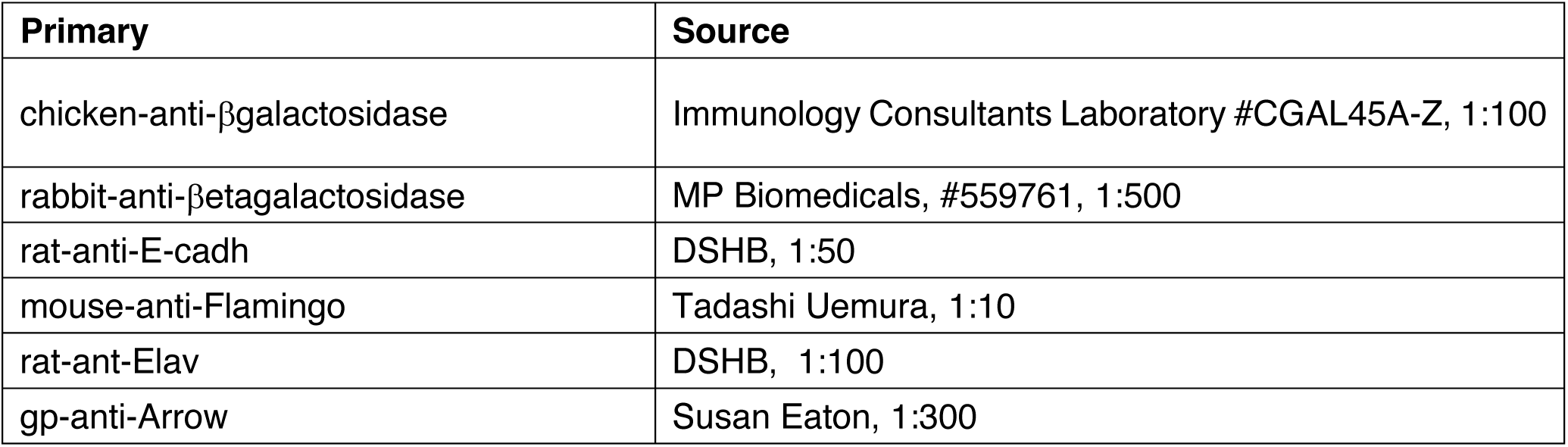
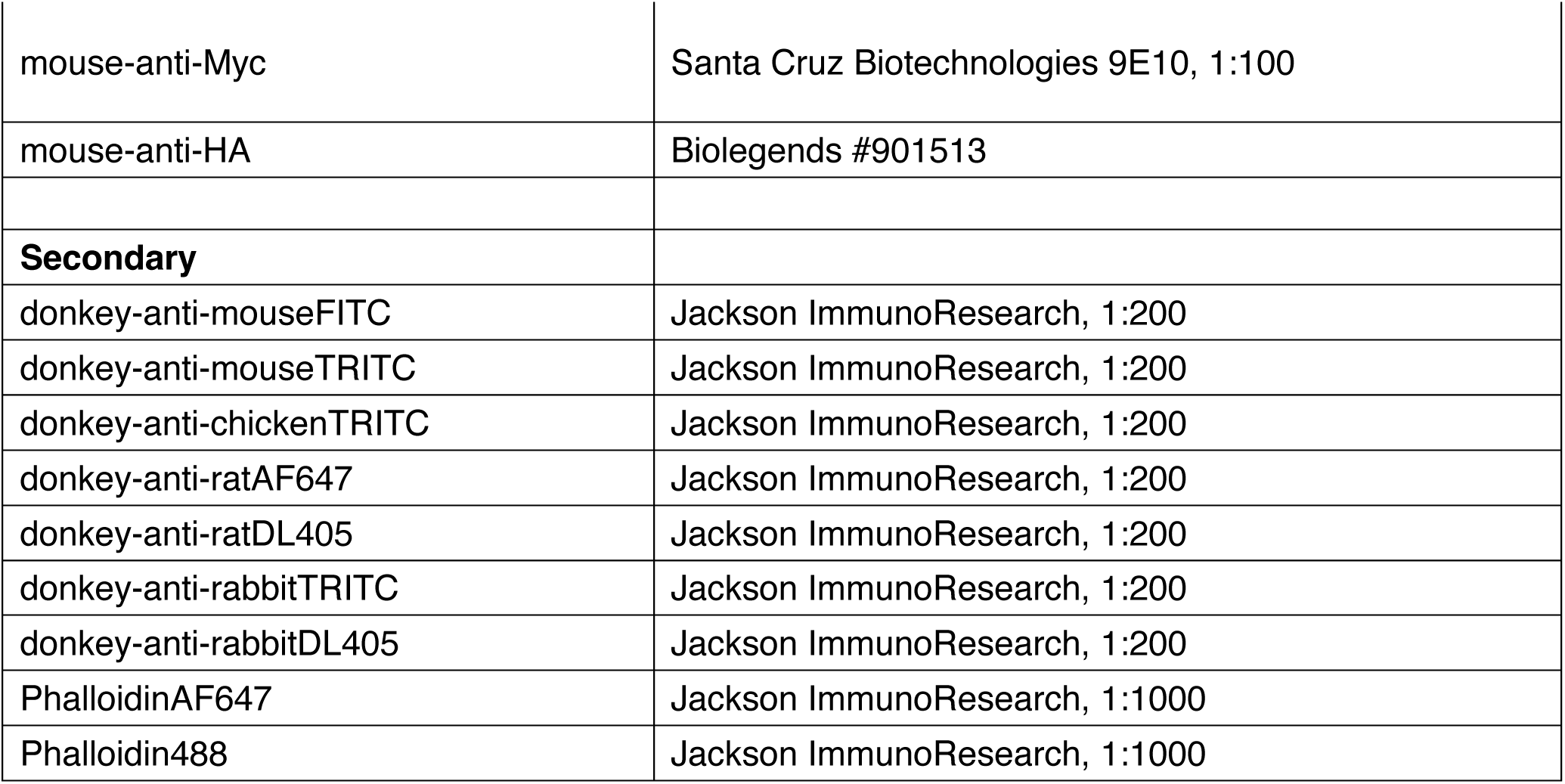

### Fluorescence image quantification and nematic order analyses

Average grey values of 18-30 cells were measured in Fiji for *dsh-DshGFP* and Arr in *hsFLP>on* Fz and Fz2 clones. Nematic order (polarity) was determined based on Fmi staining in Packing Analyzer 2.0 from 7 or 8 clones of 4 or 6 wings. Clonal and *wild-type* tissue data was extracted for further processing, omitting first row of cells in clone and wt tissue as they polarized towards clone/*wild-type* boundary.

In short, the cell segmentation information from Packing Analyzer was used to calculate the total amount of Bgal in each cell and use a threshold to distinguish mutant from *wild-type* cells. Cell assignment was manually checked to confirm correct cell type definition. The cells were then allocated into four groups: (*wt)* as *wild-type* cells that have only *wild-type* neighbors, (*wt\**) as *wild-type* cells that have at least one mutant neighbor, (*arr*^2^) and (*arr^k^*) as mutant cells that have only mutant neighbors, and (*arr*^2^*\**) and (*arr^k*^*) as mutant cells that have at least one *wild-type* neighbor.

For each cell, Packing Analyzer provides the two components of cell polarity nematic, (*p*_1_, *p*_2_). From this, we calculate polarity strength, 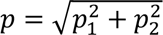, and the nematic angle, which is between −90° to 90°. Figure 2E shows violin plot distribution of *p* for pooled *wild-type* cells, *arr*^2^, and *arr^k^* mutant cells. The p-value is calculated using the two-tail T-test in Excel.

To calculate the distribution of the nematic angle, we first subtract the angle of each cell from average angle of *wild-type* cells in the respective image, allowing to pool data from several pupal wing experiments. The nematic angle distribution for all *wild-type, arr*^2^, and *arr^k^* mutant cells is shown in Figure 2F as rosette plots, and for each group of cells in Supplementary Figure S2E. To statistically compare the variance in the nematic angle distribution between different group of cells we used the F-test in Excel.

### Rh1GFP imaging

Red eyed flies give best imaging contrast. Anesthesized flies were immobilized on liquid 1.5 % agarose in a petridish at 55°C, oriented laying on their side wings in contact with agarose surface, put on ice until agaorose gelled and covered with cold distilled water, 0.01% triton X-100. Flies were kept on ice. Heads were adjusted to be able to take concentric images of the eye with a 40x water immersion lens under UV illumination. Images at different focal depths were recorded with a Zeiss camera and ommatidial phenotypes scored manually through the stack.

