## Supplemental Information for "The Wnt co-receptor Arrow-LRP5/6 is required for Planar Cell Polarity establishment in *Drosophila*"

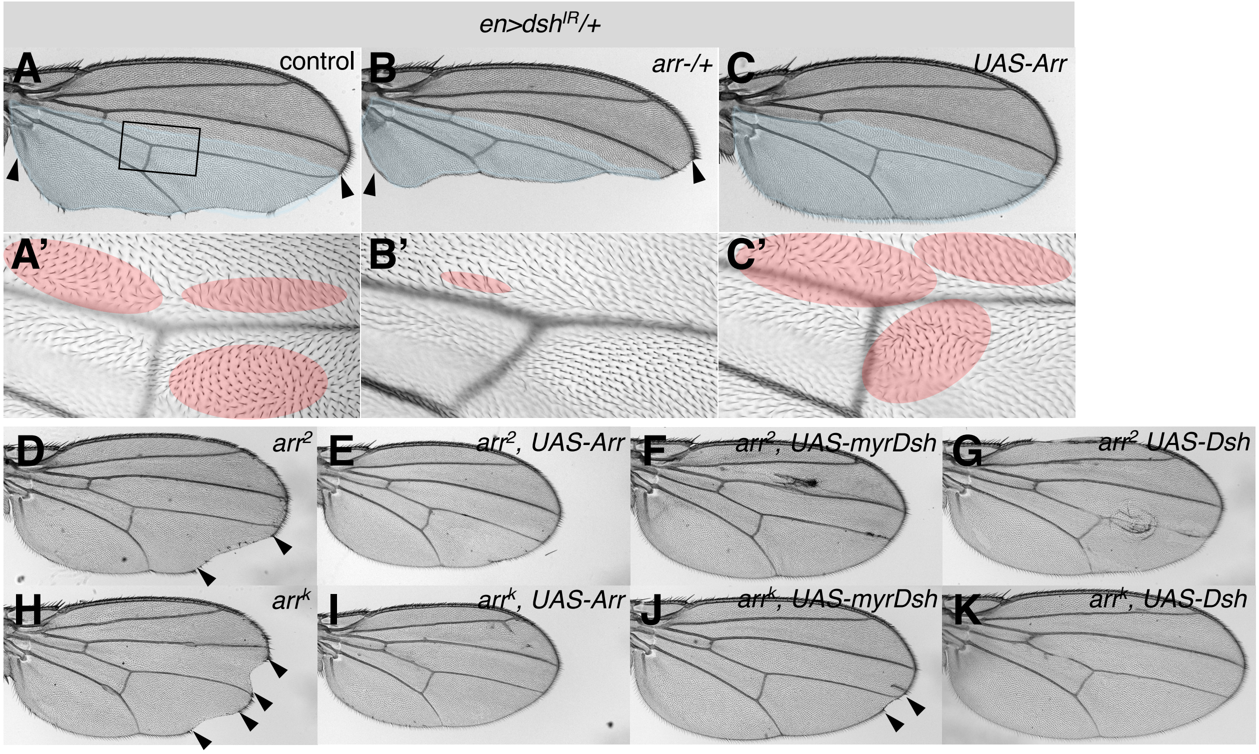

**L**

| Genotype | Wings with clones (n) | Notched wings (n) | % Notched wings |
|---|---|---|---|
| *arr[2]* | 39 | 26 | 66.7% |
| *arr[2], UAS-Arr* | 35 | 4 | 11.4% |
| *arr[2], UAS-myrDsh* | 71 | 1 | 1.4% |
| *arr[2], UAS-Dsh* | 82 | 7 | 8.5% |
| *arr[k]* | 65 | 31 | 47.7% |
| *arr[k], UAS-Arr* | 23 | 10 | 43.5% |
| *arr[k], UAS-myrDsh* | 26 | 15 | 57.7% |
| *arr[k], UASDsh* | 35 | 6 | 17.1% |
| *ALZ, UAS-myrDsh* | 22 | 1 | 4.5% |
| *ALZ, UAS-Dsh* | 47 | 3 | 6.4% |

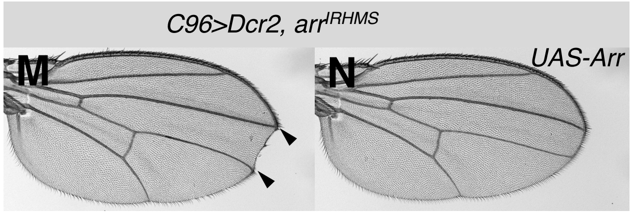

**O**

| Genotype | Total wings (n) | % Notched wings |
|---|---|---|
| *arr-IRHMS* | 29 | 58.6% |
| *arr-IRHMS, UAS-Arr* | 71 | 7.0% |

**Figure S1. Changing *Arrow* levels shifts *dsh* knockdown induced phenotypes from Wnt/βcatenin to Wnt/PCP, and rescue of *arr* mutant defects.**

(A-C) Wings with *dsh* knockdown in posterior compartment (*en-Gal4*) highlighted in blue. Genotypes as indicated, control wings were *arm-/+, nkd-/+* or *ry$^{506}$-/+*. Extent of margin loss is indicated by arrowheads. (A'-C') Area at posterior crossvein, boxed in (A), showing unaffected tissue in upper part and trichome misorientations in knockdown area highlighted in red. Reduction of *arr* copy number deepens and extends notching while suppressing trichome orientation defects (B,B'), *Arr* overexpression caused the opposite effects and ectopic margin bristles (C,C').

(D-K) Wings containing clones and clonal rescue as indicated. Control is *arm-lacZ* (*ALZ*) in (L). *arr*$^{2}$ and *arr*$^{k}$ clones induce notches (indicated by arrowheads), ectopic margin bristles, and trichome orientation defects (D,H, main fig. 1B',B",C'). *Arr*, *myrDsh* and *Dsh* rescue margin defects (E-G,I-K,L). Table summarizes extent of rescue (L). *arr*$^{2}$ is rescued in all assays with reduced occurrence of notches, *arr*$^{k}$ was rescued with *UAS-Dsh*, other rescues only reduced notch size. (M-O) *arr* knockdown in wing margin via *C96-GAL4* induces loss of wing margin (notching), indicated by arrowheads (M). Arr overexpression rescues notching phenotype (N,O), validating *IR* and overexpression tools.

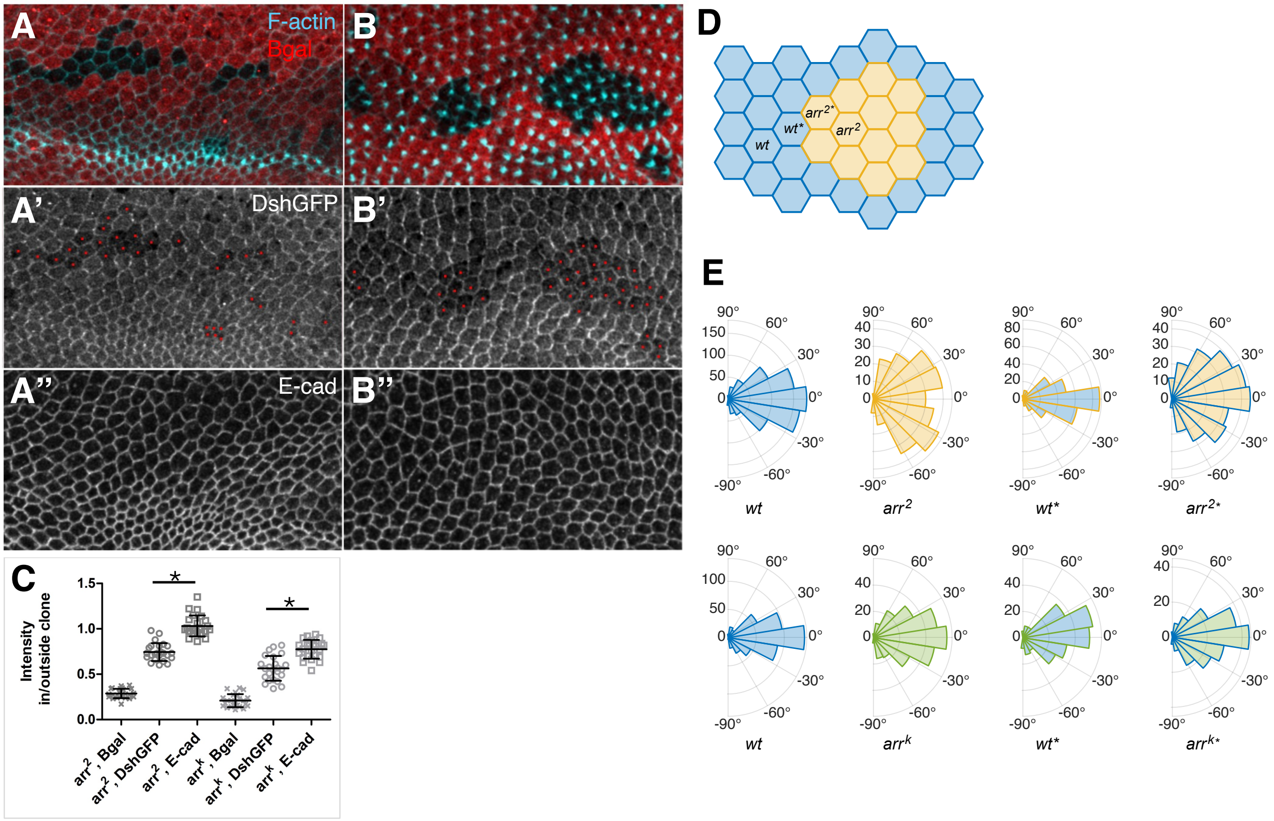

**Figure S2. *arr* LOF cells show reduced DshGFP levels with loss of polarized localization.**

(A,B) DshGFP staining in *arr²* clones, marked by loss of βgal (in red), and F-actin (in cyan). Note reduced DshGFP levels (monochrome in A',B'). (C) Quantification of DshGFP levels relative to controls for both alleles. (A-A'') Prior to prehair formation, DshGFP is reduced and asymmetric localization is lost in *arr* mutant cells, indicated by red dots (A'). DshGFP accumulates at membranes shared between mutant and *wild-type* cells (A',B'). (B-B'') At time of prehair formation, DshGFP asymmetric localization and levels are also reduced in *arr* mutant cells (B'). E-cad levels are not affected in clones (A'',B''). (C) Quantification of staining intensities in mutant versus *wild-type* cells, indicated as relative intensity in/outside clone for clonal marker βgal, DshGFP (which is reduced), and E-cad as control, from 6-9 wings and 24 clones total. T-test, * P<0.0001 for DshGFP compared to E-cad for both alleles.

(D) Schematic displaying the distinct cell distribution in which the cellular nematic order was analyzed. (E) Rosette plot quantifications of angle distribution in *arr²* (upper graphs) and *arrᵏ* mutant cells (lower graphs) and the respective cells at clonal boundaries. Note that angle distribution is randomized in mutant cells as compared to *wt* control tissue, and also that *wt* cells touching clonal borders (*wt**) can display different distributions than their respective *wt* neighbors (Fig. 1). Statistical analyses of angle distributions were calculated with F-test: *wt* vs arr2, p=2.655E-07; *wt* vs *arr²**, p=4.564E-07; *wt* vs *wt**, p=0.0233; *arr²* vs *arr²**, p=0.895; *wt* vs *arrᵏ* p=0.0339; *wt* vs *arrᵏ** p=0.0183; *wt* vs *wt** p=0.0121; *arrᵏ* vs *arrᵏ** p=0.699. These quantifications are consistent with the observation that the *arrᵏ* allele is a hypomorph, displaying weaker defects than *arr²*.

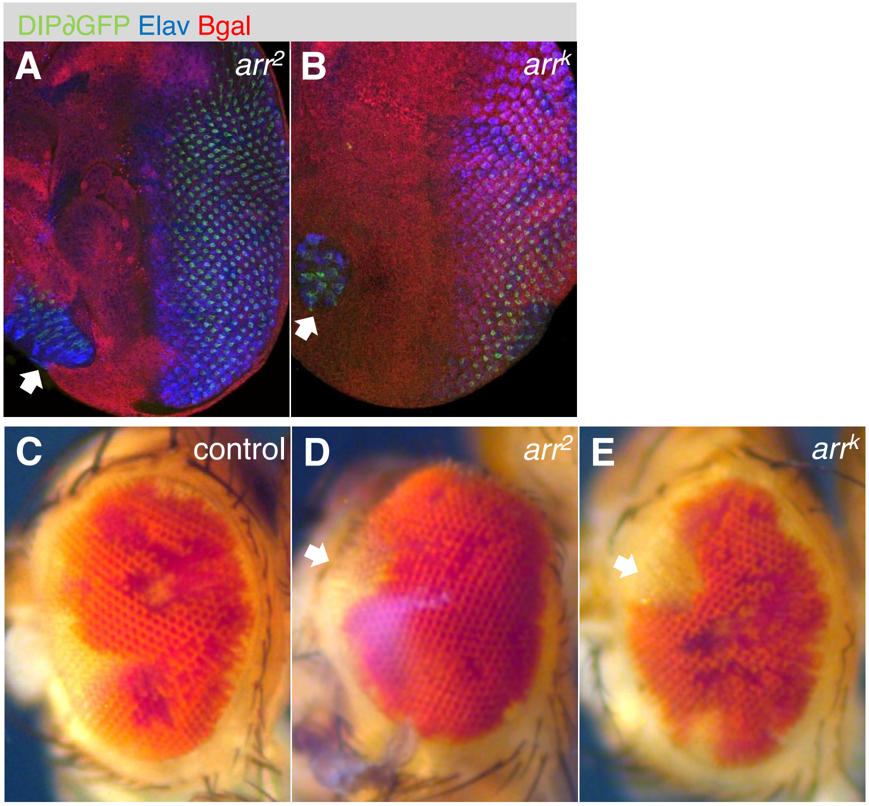

**Figure S3. *arr* clones induce ectopic eye fields in polar and anterior region leading to misshapen eyes.**

Developing (A,B) and adult eyes (C-E) with *arr* mutant tissue, genotypes as indicated. (A,B) Clones marked by loss of βgal (red), photoreceptors labelled by Elav (blue) and R3 fate (*DIPδGFP*, green). Both *arr* alleles induce ectopic eye fields at the poles and anterior to the morphogenetic furrow, examples marked by white arrows (A,B). Such ectopic eye fields cause bulging areas in adult eyes, as eye fields merge during development (examples indicated by white arrows) (D,E). (C) Control clones in adult (marked by loss of pigment marker, *white*).

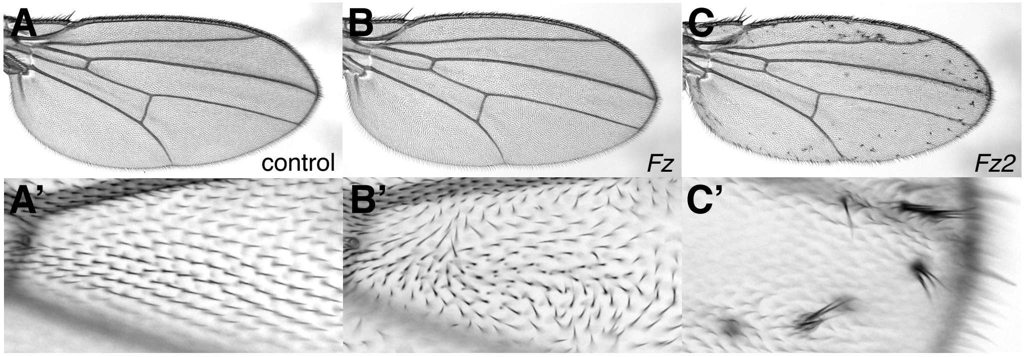

**Figure S4. Fz or Fz2 overexpression induces Wnt/PCP or Wnt/β-catenin signaling defects, respectively.**

(A-C) Adult wings with overexpression clones of *lacZ* (control)*, Fz or Fz2* induced via *hsFLP; act>y+>GAL4, UAS-GFP* from crosses used in Fig. 4. (A’-C’) Enlarged view of anterior crossvein area or tip of wing. (A,A’) Control wing (*lacZ*) with wild-type trichome orientation. Fz overexpression induces trichome misorientations and multiple trichomes (Wnt/PCP defects) (B,B’), and Fz2 induces ectopic margin bristles in wing blade (Wnt/β–catenin defects) (C,C’), confirming their known activities in these assays.

| Genetic manipulation | Wnt/PCP signaling phenotype | Wnt/β−catenin signaling phenotype | Assay |
|---|---|---|---|
| *dsh* k.d. | +++ | +++ | *en>dshIRT* |
| *dsh* k.d., less *arr* | + | +++++ | wing |
| *dsh* k.d., more *arr* | +++++ | + | Sup. fig. 1 |
| Dsh o.e. | +++ | +++ | *sep>Dsh* |
| Dsh o.e., less *arr* | + | + | eye |
| Dsh o.e., more *arr* | +++++ | +++++ | Fig. 5 |
| Fz o.e. | +++ | n.a. | *sev>, pnr>Fz* |
| Fz o.e., less *arr* | n.e./+ | n.a. | eye/notum |
| Fz o.e., more *arr* | +++++ | n.a. | Sup. fig. 5/ Fig. 5 |
| *arr* k.d. | n.e./* | +++ | *en>arr-IR* |
| *arr* k.d., less *fz* | n.d. | + | wing |
| *arr* k.d., more *fz* | n.d. | +++++ | Sup. fig. 5 |
| *arr* LOF | +++ | +++ | MARCM clones |
| *arr* LOF, more (myr)dsh | + | + | wing, Sup. fig. 1 |

**Supplemental Table 1. Summary of functional interactions between *arr*, *fz* and *dsh* in Wnt/PCP and Wnt/β−catenin pathways**.

Left column lists genetic manipulations of *dsh*, *fz* or *arr* via knockdown (k.d.), overexpression (o.e.) or LOF clones (LOF) and decrease (less) or increase (more) of *arr*, *fz* or *dsh* via knockdown, overexpression or copy number reduction. Second and third columns indicate phenotypic effects caused by manipulations and how their relative strength changes, marked by + for either signaling pathway, indicated in header. Fourth column lists genotypes, which tissue was examined and reference to specific figures. No effects (n.e.), * = *arr* clones induce PCP defects, for full genotypes see Methods.

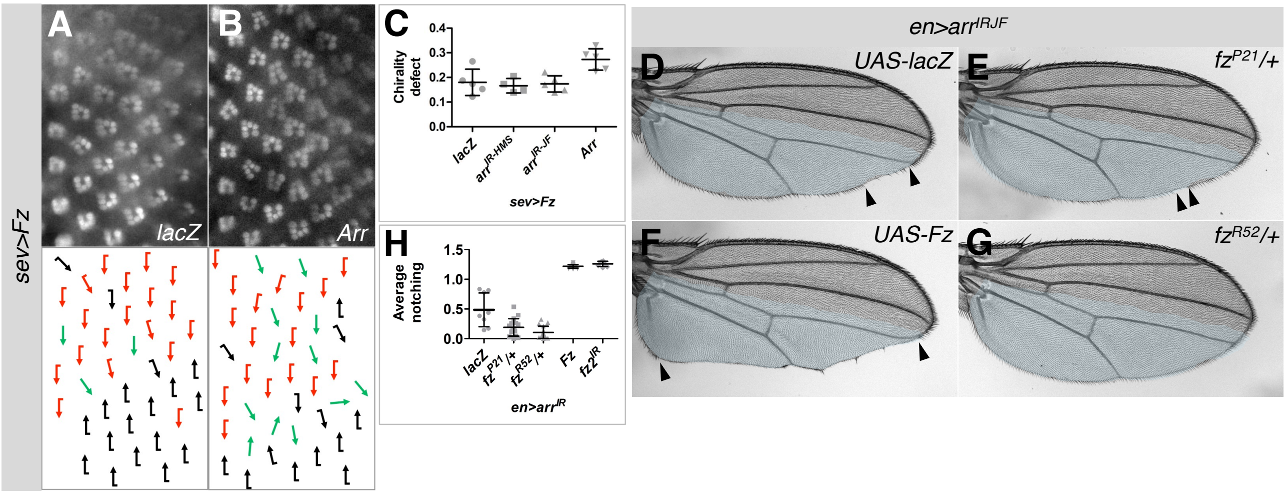

**Figure S5. Fz induced PCP defects are enhanced by Arr co-expression, and Fz titrates Arr away from β-cat signaling.**

(A,B) Top panels show rhabdomere arrangement, visualized with *Rh1GFP* (anterior is left) in *sev>Fz* (for precise genotypes see Methods). Genotypes as indicated. Schematic in lower panels with red and black arrows representing dorsal or ventral chirality, respectively. Green arrows designate ommatidia with symmetric arrangement of the R3-R3 type. *sev>Fz* induces inversed chirality (intermixed red and black arrows) and R3-R3 symmetric ommatidia (A-C, compare to *wild-type* in Fig. 5A.). Arr overexpression enhances chirality defects (B,C). (C) Quantification summarizing ommatidial chirality/PCP defects. *arr* knockdown did not modify Fz overexpression significantly. Knockdown or overexpression of *arr* alone did not induce phenotypes. t-test: *P*=0.0169 for enhanced chirality defects compared to control (*lacZ*), 4-5 eyes each, n=305-694 ommatidia analyzed.

(D-G) Fz titrates Arr away from β-cat signaling. Adult wings with *arr* knockdown in posterior compartment (highlighted in blue) display margin loss/notches, extent marked by arrowheads (for precise genotypes see Methods). Reducing *fz* copy number dominantly suppresses this phenotype (E,G,H), while Fz overexpression enhances it comparably to a *fz2* knockdown (F,H). (H) Quantification of relative margin loss (notching) compared to length of L3 in *en>arr$^{IR}$* wings. t-test: *P*<0.0001 for all compared to *lacZ*, 16 wings each

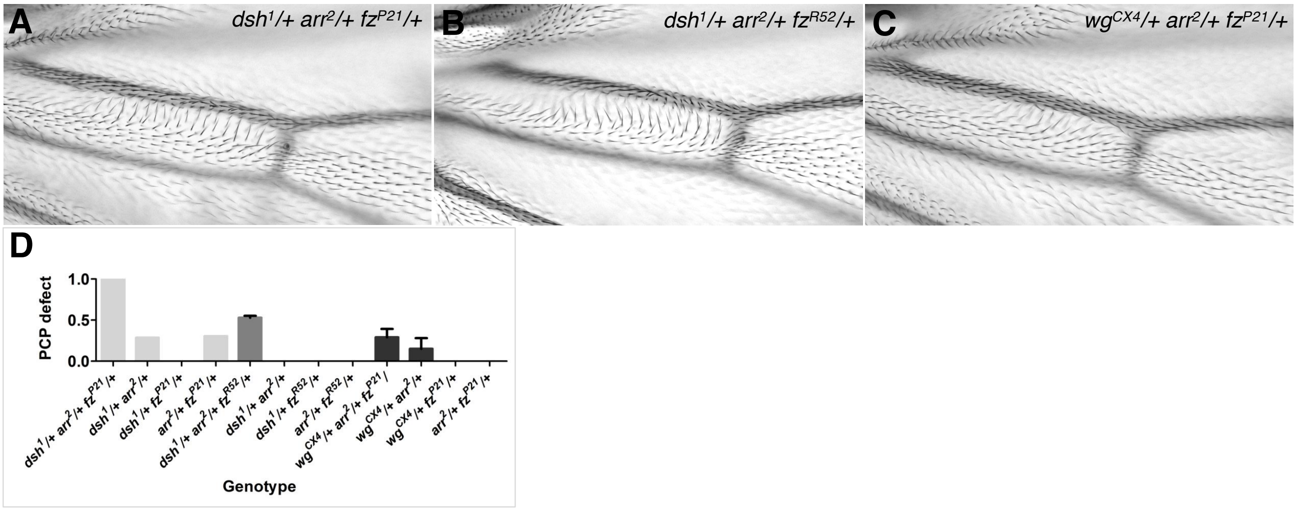

**Figure S6.** ***wg, arr*, *fz*, or *dsh* multi-heterozygous conditions result in PCP defects.**

(A-C) Wings multiply heterozygous for *dsh*, *arr*, *fz* or *wg*, with focus on the *dsh¹* allele. Genotypes as indicated. Proximal area to anterior crossvein between vein L3 and L4 is shown, which is sensitive to PCP perturbations. Note trichome misorientations (A-C) compared to control wings (main Fig. 6A). Quantification summarizing % of PCP defects observed (D). Triple heterozygous *dsh-/+, arr-/+, fz-/+* wings show 100% PCP defect (A,D), compared to *dsh-/+, arr-/+* (0%) or *arr-/+, fz-/+* (0%) and *dsh-/+, fz-/+* (30%). Similar effects were seen for an independent *fz* allele, triple hetrerozygous wings show 66.7% PCP defect (B,D), compared to other double heterozygotes (0%). n=14-54 wings each. Note that combinations with the hypomorphic allele $dsh^1$ yield generally a lower phenotypic frequency than the *dsh* null allele, $dsh^{V26}$ (Fig. 6). Triple heterozygous *wg-/+, arr-/+, fz-/+* wings show 29.1% PCP defect compared to *wg-/+, arr-/+* (15%), and others 0% defects. Note, the single *wg-/+* mutant combination phenotypes are less penetrant than the double mutant *wg*, *Wnt4* defects shown in Fig. 6.
